# A dual component system instructs membrane hydrolysis during the final stages of plant autophagy

**DOI:** 10.1101/2025.08.01.668046

**Authors:** Julie Castets, Matthieu Buridan, Inés Toboso Moreno, Víctor Sánchez de Medina Hernández, Rodrigo Enrique Gomez, Franziska Dittrich-Domergue, Josselin Lupette, Clément Chambaud, Stephanie Pascal, Tarhan Ibrahim, Tolga O. Bozkurt, Yasin Dagdas, Frederic Domergue, Jérôme Joubès, Elena A. Minina, Amélie Bernard

**Affiliations:** Univ. Bordeaux, CNRS, LBM, UMR 5200, Villenave d’Ornon, F-33140, France; Gregor Mendel Institute (GMI), Austrian Academy of Sciences, Vienna BioCenter (VBC), Vienna, Austria; Vienna BioCenter PhD Program, Doctoral School of the University of Vienna and Medical University of Vienna, A-1030, Vienna, Austria; Department of Life Sciences, Imperial College London, London, United Kingdom; Heidelberg University, Centre for Organismal Studies (COS), 69120 Heidelberg, Germany; Department of Molecular Sciences, Uppsala BioCenter, Swedish University of Agricultural Sciences and Linnean Center for Plant Biology, PO Box 7015, SE-75007 Uppsala, Sweden

**Keywords:** Autophagy, autophagosomes, autophagic bodies, phospholipase, membrane, *Arabidopsis*

## Abstract

Autophagy is an intracellular catabolic process conserved across eukaryotes and critical for plant stress tolerance. Upon their delivery in the vacuole, how autophagic bodies containing cargo are hydrolyzed to warrant autophagy degradation remains poorly characterized. Here, we identify two Arabidopsis phospholipases as core components of the autophagy machinery. We find that LCAT3 and LCAT4 traffic to the vacuolar lumen and converge on autophagic bodies using differential pathways, placing them on the outer and inner side of the vesicle, respectively. Double knockouts *lcat3,4* accumulate autophagic bodies and show reduced autophagy activity. *In vivo* reconstitution demonstrates that LCAT3 can hydrolyze the membrane of autophagic bodies, enabling the activity of LCAT4 to enhance this process. In sum, our work reveals that the concerted action of a multi-component system is required for the efficient and specific disruption of autophagic bodies as an obligatory step for the completion of the autophagy pathway.

## Introduction

Eukaryotes have developed macroautophagy (hereafter referred to as autophagy), an intracellular catabolic pathway which integrates environmental and developmental cues to promote cell homeostasis and adaptation. This process entails the sequestration of intracellular cargo (proteins, protein aggregates, organelles) into specialized vesicles, its trafficking to the lytic compartment of the cell (the lysosomes in mammals and the central vacuole in yeast or plant cells), its degradation therein and the subsequent recycling of the resulting molecules (Gomez et al., 2021). The importance of autophagy in plant physiology is unequivocal: it is critical for plant survival to virtually all types of stresses and, although plants lacking autophagy can grow in controlled conditions, they show pleiotropic developmental phenotypes including accelerated senescence, defects in seed formation and germination as well as reduced vegetative growth and fecundity (Gross et al., 2025). The physiological importance of autophagy is mediated by its two main functions: the degradation of damaged, unwanted or superfluous intracellular material which both detoxifies cells and rewires their activities to meet developmental and environmental demands, and the recycling of autophagy end-products, which reallocates resources when they become scarce and sustains metabolism thereby supporting cell survival (Gomez et al., 2021). As such, autophagy is defined as a degradation and recycling process.

At present our understanding of autophagy has largely focused on the initial stages of the pathway, *i.e.*, the formation of autophagy vesicles, named autophagosomes, and the molecular bases of cargo recognition and sequestration (Zhuang et al., 2024; Otegui et al., 2024; Gross et al., 2025). Upon induction, autophagy morphologically starts with the nucleation of a cup-shaped membrane compartment, the phagophore, at which the autophagy machinery sequentially coalesces. This machinery consists in a set of well-characterized proteins named autophagy related proteins (ATG proteins), initially discovered in yeast and largely conserved in mammals and plants. Among those, ATG8 is a central actor of the pathway, which recruitment at the phagophore participates in cargo selection through protein/protein interactions (Zhuang et al., 2024; Otegui et al., 2024). The sequential action of the ATG machinery results in the structurally- and functionally-hyper organized expansion of the phagophore, which progressively grows while recognizing and engulfing autophagy cargo until the fusion of the membranes at the rim of the phagophore which yields a double membrane autophagosome packed with material to be degraded (Zhuang et al., 2024; Otegui et al., 2024; Gomez et al., 2018). Either directly or upon maturation with late endosomes or multivesicular bodies (MVB), autophagosomes ultimately fuse with the vacuolar membrane (Zhao et al., 2022; Jiang et al., 2024). At this step, the outer membrane of autophagosomes/amphisomes becomes part of the tonoplast, while their inner membrane containing cargo, are released as a vesicle named autophagic body in the vacuolar lumen (Marshall and Vierstra 2008; Zhuang et al., 2024).

Little is known about the fate of autophagic bodies; yet, their breakdown represents a pivotal entry point into vacuolar degradation as it releases cargo for subsequent hydrolysis and thereby controls the completion of the autophagy pathway and its function as a degradation mechanism. More than a mere passive event, the disruption of the membrane of autophagic bodies must be tightly controlled in time, to process the large influx of vesicles in the vacuole upon autophagy induction, and in space, to localize membrane disruption at autophagic bodies while preserving the homeostasis of the vacuolar membrane. In particular, the plant lytic vacuole shows distinct features compared to the small and highly mobile lysosomes or to the yeast vacuoles where some aspects of autophagy degradation have been characterized (Kagohashi et al., 2023). The vacuole represents up to 90% of the whole volume of plant cells (Marty, 1999); its membrane, the tonoplast, compartmentalizes an acidic sap (pH ∼5-5.5; Shen et al., 2013) where hydrolases perform the lytic functions of the cell; consequently, rupture of the tonoplast leads to rapid cell death (van Doorn et al., 2011). Contrarily to yeast or mammals where the membrane of lytic compartments is protected from intralumenal hydrolases by the high glycosylation of resident proteins (forming the so-called glycocalyx; Frain et al., 2024), plant vacuolar proteins show little glycosylation (Pedrazzini et al., 2016) questioning the presence of a protective glycocalyx in plant cells. At this time, how the plant vacuole processes the large influx of autophagic bodies observed upon autophagy induction (Gomez et al., 2021), and their specific hydrolysis to complete autophagy degradation while maintaining the integrity of its membrane remains completely unknown.

Our proteomic analyses identified an atypical phospholipase A (PLA) called LCAT4 enriched in purified autophagy compartments. Phospholipases A are enzymes which catalyze the hydrolysis of acyl chains from phospholipids, contributing to lipid processing and remodeling, and thereby regulating membrane composition, function, stability and biophysical properties (Zhang et al., 2021). As such, phospholipases A are involved in numerous developmental and cellular processes including the morphogenesis of endomembrane compartments as well as plant responses to stresses (Takáč et al., 2019). However, none of these PLA has yet been linked to autophagy in plants and while the biochemical activity of LCAT4 has been partially characterized (Chen et al., 2012), its biological function remains unknown. At the starting point of this study, we aimed at addressing the physiological relevance of the association of LCAT4 with autophagy compartments, *i.e.*, the potential implication and function of LCAT4 in the autophagy pathway. In this work, combining cell biology, molecular modeling, biochemistry, genetics and *in vivo* reconstitution, we show that LCAT4 uses autophagy to relocalize from the cytosol to the vacuole inside autophagic bodies upon nutrient starvation. We found that the closest homolog of LCAT4, LCAT3, traffics to the vacuole independently of autophagy, where it recognizes the outer leaflet of the membrane of autophagic bodies. In the absence of both enzymes, autophagic bodies accumulate in the vacuole and autophagic flux is largely reduced. Conversely, expressing a combination of LCAT3 and LCAT4 in yeast cells efficiently disrupts the membrane of autophagic bodies. Together, this study leads us to propose a model in which LCAT3 and LCAT4 act synergistically to support the specific and efficient disruption of autophagic bodies in the vacuole of *Arabidopsis*, thereby unraveling the mechanism and molecular actors involved in the vacuolar stage of autophagy degradation.

## Results

### LCAT4 localizes to autophagy compartments

During autophagy, the formation and subsequent disruption of autophagosomes rely on extensive membrane remodeling events. Lipids and proteins involved in lipid dynamics play critical roles in the instruction of membrane shaping. In that context, and to better understand the molecular bases underlying autophagy, we searched for proteins involved in lipid metabolism in the proteome of autophagy compartments that we pulled down with ATG8A, a core component of the autophagy machinery. Alongside other autophagy-related proteins, we found that a phospholipase A called LCAT4 is enriched in GFP-ATG8A decorated membranes (**Fig. 1a**), suggesting that this soluble protein localizes in autophagy compartments. To get insights into a possible link between LCAT4 and autophagy, we generated stable Arabidopsis lines co-expressing LCAT4-RFP and GFP-ATG8A, to analyze their respective subcellular localization. Using *in vivo* confocal microscopy of *Arabidopsis* seedlings roots, we observed that under nutrient-rich conditions (+NC), LCAT4 is mostly found in the cytosol and on puncta co-localizing with ATG8A (**Fig. 1b-d**). When autophagy is induced, ATG8A is recruited to the phagophore membrane and a portion of the protein remains associated with completed autophagosomes; as such, part of the protein is delivered into the vacuole and found inside autophagic bodies (Zhuang et al., 2024). As expected, when autophagy was induced after 1h of nutrient deprivation (lack of nitrogen and carbon, -NC), the number of GFP-ATG8A-labeled puncta increased (**Fig. 1b,c**). Consistently, we observed an increase in the number of LCAT4 puncta **(Fig. 1b-c**) with a high percentage of colocalization with GFP-ATG8A (**Fig. 1d**). To observe autophagic bodies, we used concanamycin A (CA) which de-acidifies the vacuole, resulting in a reduction of vacuolar degradation (Huss et al., 2002). In this condition, we observed a large accumulation of LCAT4-RFP puncta co-localizing with GFP-ATG8A inside the vacuolar lumen (**Fig. 1c-d**). Similarly, in lines expressing LCAT4-BFP and YFP-ATG18A, a phagophore marker, BFP signal was found on ring-like structures colocalizing with ATG18A (**Fig. S1**). These results show that LCAT4 is associated with all autophagy compartments, *i.e.*, phagophores, autophagosomes and autophagic bodies.

**Figure 1.**
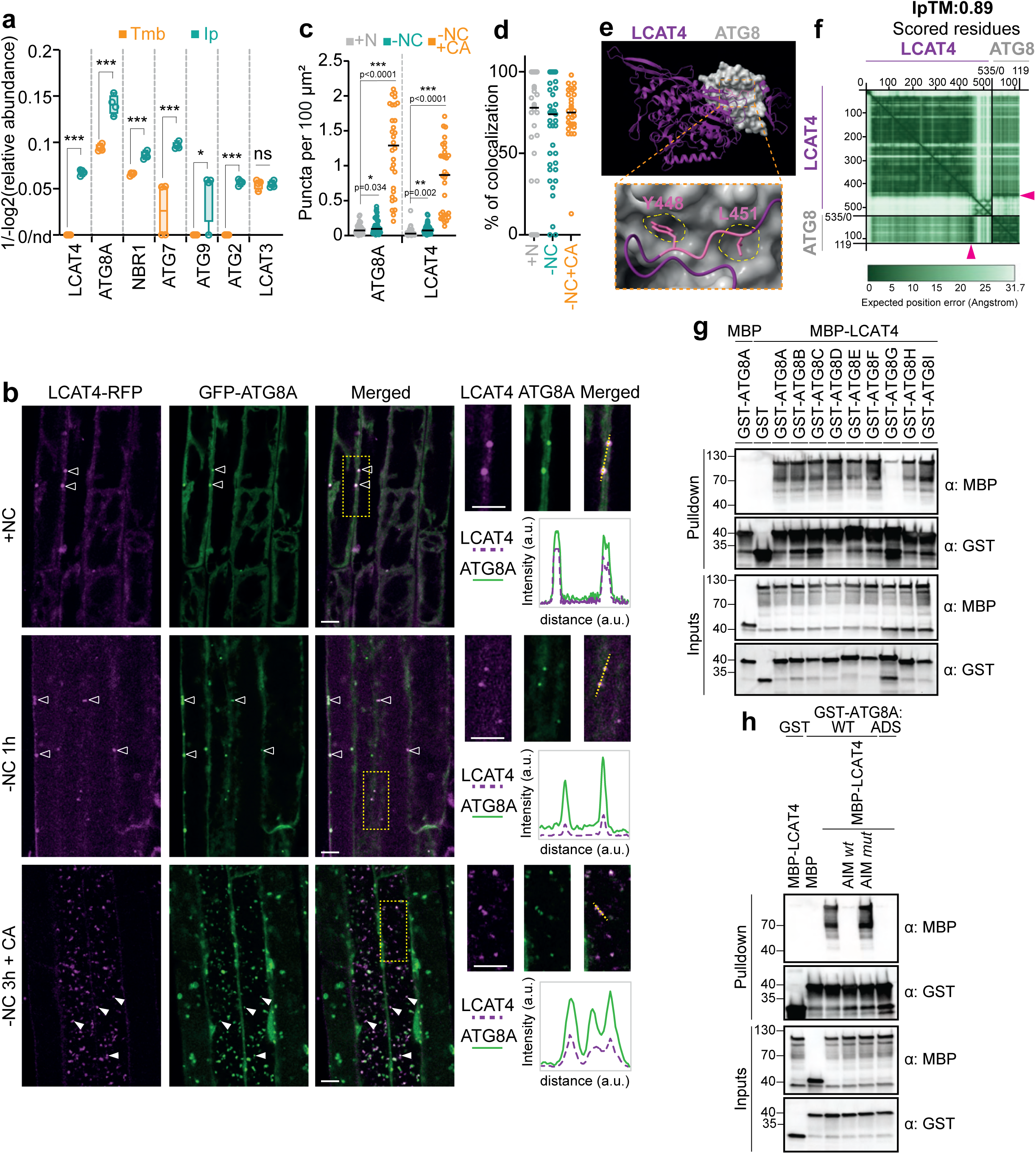
LCAT4 localizes to autophagy compartments and directly interacts with ATG8A. **(a)** LCAT4 is enriched in immuno-isolated ATG8A compartments, similarly to resident markers of the autophagy machinery (NBR1, ATG7, ATG9, ATG2) while LCAT3 is not. Results represents the 1/-log2 of relative abundance of each protein in ATG8A-compartments (Ip) compared to the membrane fraction input (Tmb). Data are presented as box plot with min to max values, average and all individual values of n=4 independent biological replicates. ns, non-significant, *, p<0.05, p<0.01 and ***, p<0,001 with student t-test. **(b)** LCAT4 localizes in the cytosol and co-localizes with ATG8A on autophagy compartments. Representative confocal images of 7-day-old roots of seedlings co-expressing LCAT4-RFP and GFP-ATG8A. Plants were placed in rich condition (+N), deprived of nutrients for 1h (-NC 1h), or in -NC supplemented with concanamycin A (1 μM) for 3 h (-NC 3h + CA). Yellow dotted rectangles are enlarged on the right. Signal intensity profiles along the dotted lines are plotted in the boxed regions below and show co-localization between ATG8A and LCAT4 puncta. Scale bar, 10 μm. **(c,d)** Quantification of the GFP-ATG8A and LCAT4-RFP signal in (b) shows an increased number of puncta upon -NC and -NC +CA (c) and a large co-localization between the two proteins in each condition (d). Co-localization is represented as the percentage of RFP puncta colocalizing with GFP puncta (d). In +N, n=40 images were examined over 12 roots in three independent biological experiments, in -NC, n=39 images were examined over 12 roots in three independent biological experiments and in -NC + CA, n=30 images were examined over 9 roots in two independent biological experiments. All individual values are presented with median, p values of student t-test are indicated. **(e,f)** AlphaFold3-multimer modeling (e) and predicted aligned error (PAE) plot (f) show a mostly organized conformation of LCAT4 and place ATG8 in close proximity to the residues Y448 V449 I450 L451 of LCAT4 (see arrow). Y448 and L451 are predicted to bind to the W and L pockets of the ATG8 ADS domain (shown as yellow dashed circles, e). **(g,h)** LCAT4 binds ATG8 in an AIM dependent manner *in vitro*. MBP-LCAT4 is pulled-down by each of the nine Arabidopsis ATG8 isoforms (g). In contrast, interaction is lost when the ADS mutant of ATG8 is used (ATG8A^ADS^=ATG8A^(Y50A,L51A)^) or when AIM competition peptides are added (AIM *wt* compared to AIM *mut* peptides, added to a final concentration of 200 µM; h). Bacterial lysates containing recombinant protein were mixed and pulled down with glutathione magnetic agarose beads. Input and bound proteins were visualized by immunoblotting with anti-GST and anti-MBP antibodies. Images are representative of n=2 independent experiments (see **Fig. S3**).

We next wondered how LCAT4 was recruited and associated to autophagy compartments. Analyses of the amino acid sequence of LCAT4 found 11 putative ATG8 interaction motifs (AIM; with a consensus site characterized as [W,Y,F][X][X][L,I,V], Ibrahim et al., 2023, see **Fig. S2a**). AlphaFold3-multimer modeling indicated a direct binding between LCAT4 and ATG8 and highlighted residues YVIL^448-451^ in the C-terminal region of LCAT4 as a potential AIM domain associating with the AIM docking site of ATG8 (ADS, **Fig. 1e-f**; **Fig. S2b-e**). The potential interaction of LCAT4 with ATG8 was thus tested using recombinant proteins and pull-down assays. These analyses showed that MBP-LCAT4 is pulled-down by each of the nine ATG8 isoforms from Arabidopsis (ATG8A-I) indicating a direct interaction between ATG8 and LCAT4 (**Fig. 1g; Fig. S3a**). Moreover, this interaction was drastically reduced when a competitive AIM peptide, binding to the ADS domain of ATG8A, was added in the reaction mixture or when we used a form of GST-ATG8A where the ADS domain was mutated (**Fig. 1h; Fig. S3b**). These results show that LCAT4 directly binds to ATG8 and suggest the implication of a canonical AIM-ADS interaction in the localization of LCAT4 to autophagy compartments.

### LCAT4 uses autophagy as a trafficking route to reach the vacuolar lumen upon nutrient starvation

Results from **Fig. 1** show that LCAT4 is cytosolic in nutrient-rich conditions and localize to autophagosomes and autophagic bodies upon nutrient starvation. Further analyses of the localization of LCAT4 revealed that the cytosolic pool of LCAT4-RFP largely decreases after prolonged period of starvation (-NC 3h), with a concomitant increase of the protein signal inside the vacuole lumen (**Fig. 2a, c**; see co-localization with the vacuolar marker BCECF in **Fig. S4**). Further, immunoblot analyses show that the level of LCAT4 is mostly unaffected in these conditions (**Fig. 2e**), suggesting that the presence of LCAT4 in the vacuole does not lead, at least initially, to its degradation. Therefore, we concluded that LCAT4 is not a substrate for autophagy degradation but rather transported from the cytosol to the vacuole upon nutrient starvation. Because we found that LCAT4 associates with autophagy compartments in these conditions (**Fig. 1, 2**), we postulated that the protein uses the autophagy pathway as a trafficking route to reach the vacuole. To test this hypothesis, we analyzed the localization of LCAT4 and ATG8A when autophagy was inhibited either pharmacologically or genetically. Upon Wortmannin (Wm) treatment, an inhibitor of the PI3 kinase complex and of autophagy (Beesabathuni et al., 2023), we observed a large reduction of ATG8A puncta compared to untreated cells (**Fig. 2a-b**), indicating that autophagosome formation is disrupted as expected. In these conditions, the number of LCAT4 puncta also decreases, supporting the genuine association of LCAT4 with autophagosomes (**Fig. 2a,b**). Further, the LCAT4-RFP signal was found mostly absent from the vacuolar lumen but, rather, restricted to the cytosol (**Fig. 2a,c**). Similarly, when LCAT4-RFP was expressed in the *atg5* KO line, where autophagosome formation is blocked (Chung et al., 2010), we did not observe any LCAT4 puncta, and the signal of the protein was found exclusively in the cytosol (**Fig. 2b-d**). Together, these results show that LCAT4 relocates from the cytosol to the vacuolar lumen upon nutrient starvation, and that autophagosome formation is required for its transport.

**Figure 2.**
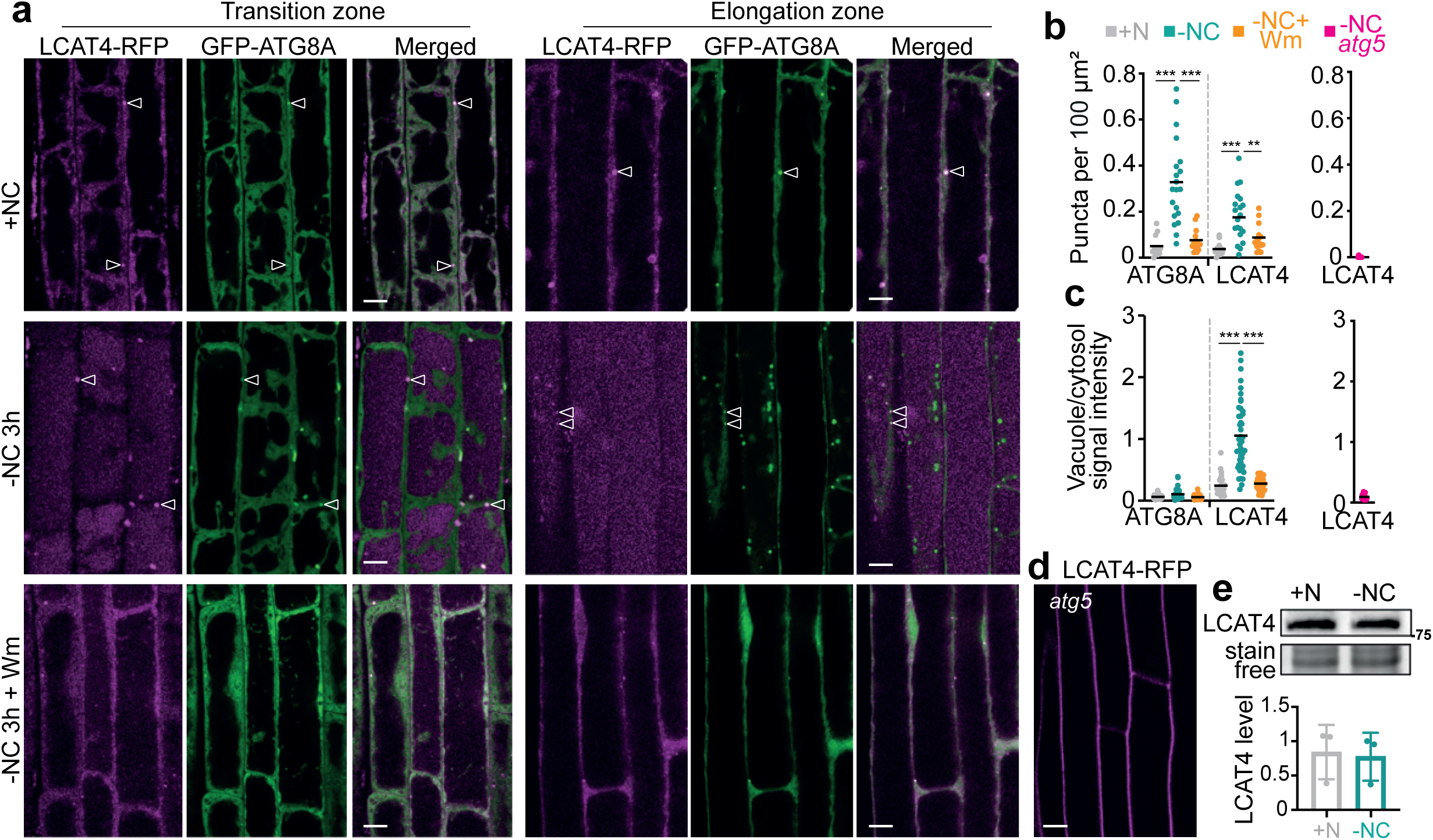
LCAT4 uses autophagy to traffic to the vacuole upon nutrient starvation. **(a-d)** LCAT4-RFP relocalizes from the cytosol to the vacuolar lumen upon nutrient starvation. Blocking autophagy (with the addition of Wortmannin or in the *atg5* mutant) restricts LCAT4-RFP in the cytosol in the same conditions. **(a)** Representative confocal images of Arabidopsis 7-day-old seedlings co-expressing GFP-ATG8A and LCAT4-RFP in the transition and elongation zone of the roots. Plants were imaged in nutrient rich condition (+N), after 3 hours in nutrient starvation (-NC 3h) or after -NC 3h supplemented with wortmannin (-NC 3h + Wm; 1 μM). Scale bar, 10 µm. **(b)** Wortmannin treatment reduces the number of ATG8A and LCAT4 puncta; similarly, no LCAT4 puncta are detected when LCAT-RFP was expressed in the *atg5* mutant. Quantification of the number of GFP-ATG8A and LCAT4-RFP puncta of images of (a, left panel) and (d, right panel). Results are presented as the number of puncta per surface of root area and show all individual values and average of: in +N, n=12, in -NC, n=20, in -NC + Wm, n=15, in *atg5*, n=5. **, p<0.01 ***, p<0,001, student t-test. **(c)** Quantification of the LCAT4-RFP signal measured in the vacuole compared to LCAT4-RFP signal measured in the cytosol of images from (a). Results are presented as the ratio of vacuole/cytosol signal intensity and show the average and individual values of: in +N, n=28 cells were examined over 8 roots in three independent biological experiments, in -NC, n=43 cells were examined over 11 roots in four independent biological experiments, in NC + Wm, n=37 cells were examined over 8 roots in three independent biological experiments, in *atg5*, n=14 cells were examined over 5 roots in one biological experiment. ***, p<0.001, student t-test. **(d)** Confocal image of 7-day-old roots of seedlings of *atg5* mutant expressing LCAT4-RFP after 3 hours in nutrient starvation. Scale bar, 10 μm. **(e)** The level of LCAT4-RFP is unaffected in -NC. Detection of LCAT4-RFP by immunoblot from total proteins extracted from 7-day-old seedlings after 3 hours in liquid rich condition (+N) or nutrient starved liquid medium (-NC). Representative image of n=3 independent experiments (upper panel). The level of LCAT4 was detected, quantified and normalized by the loading control; all values are presented with average and SD (lower panel).

### LCAT4 is dispensable for autophagy during nitrogen and carbon deprivation

We next sought to understand the functional relevance of the association of LCAT4 with the autophagy pathway. Although the function of LCAT4 remained unknown in plants, its biochemical activity was partially characterized by expressing the protein in *S. cerevisiae* (Chen et al., 2012). In this system, LCAT4 was found to have a phospholipase A_2_ activity, *i.e.*, to hydrolyze the acyl chain at the *sn-2* position of a variety of phospholipids. Further, LCAT4 was found to only be active at acidic pH, between 4.5 and 5.5, with an optimal at pH=5.0 (Chen et al., 2012), which corresponds to the pH of the vacuolar lumen of *Arabidopsis* (Shen et al., 2013). Based on our results presented in **Fig.1-2**, we thus proposed that, mostly residing in the pH-neutral cytosol, LCAT4 is inactive at steady state; upon nutrient starvation, autophagy is induced, transporting LCAT4 into the acidic vacuole where it becomes active. Because of its biochemical activity as a phospholipase, its presence in autophagic bodies and its trafficking inside the vacuole upon autophagy induction, we reasoned that LCAT4 may play a role in the hydrolysis of autophagic bodies in the vacuole during nutrient starvation. To test this hypothesis, we measured the impact of misexpressing *LCAT4* on autophagy, using a T-DNA knocked-out line, *lcat4* (**Fig. S5a-c)** as well as a line in which the expression of *LCAT4* was conditionally knocked-down (*amiRNA:LCAT4*, **Fig. S5d**). First, we measured the rates of autophagic flux using the western-blot based GFP-ATG8 processing assay. GFP-ATG8 is associated with autophagosomal membrane and a portion of the protein is delivered in the vacuole inside autophagic bodies. Upon the disruption of the membrane of autophagic bodies, GFP-ATG8 is delivered in the vacuolar lumen where it is rapidly degraded, while the more stable GFP moiety accumulates (Shin et al., 2014). Therefore, measuring the ratio between the level of GFP-ATG8A and that of its degradation product, free GFP, quantitatively reflects the rate of autophagic degradation. To test whether LCAT4 was required for autophagy activity, seedlings of *LCAT4* KO and KD lines were placed in nutrient rich liquid medium (+N) or nutrient-depleted liquid medium (lacking either nitrogen, -N or a combination of nitrogen and carbon, -NC) and compared to WT plants. Upon -N or -NC, autophagy was induced in WT plants, as seen by the reduction of the GFP-ATG8A signal and the concomitant increase in free GFP (**Fig. 3a,b**). The ratio of GFP/GFP-ATG8A was not significantly different than that of WT in plants KO or KD for *LCAT4* (**Fig. 3a,b** and **Fig. S6,** respectively) indicating that autophagy flux is not affected by the absence of LCAT4. Similarly, *in vivo* quantitative analyses of the number of autophagosomes (**Fig. 3c,d**) and autophagic bodies (**Fig. 3c,e**) labelled by GFP-ATG8A in the roots of 7-day-old seedlings showed no significant differences between WT and *lcat4* plants. Together, these results show that the absence of LCAT4 does not significantly impact autophagosome formation, autophagic body accumulation or autophagic degradation.

**Figure 3.**
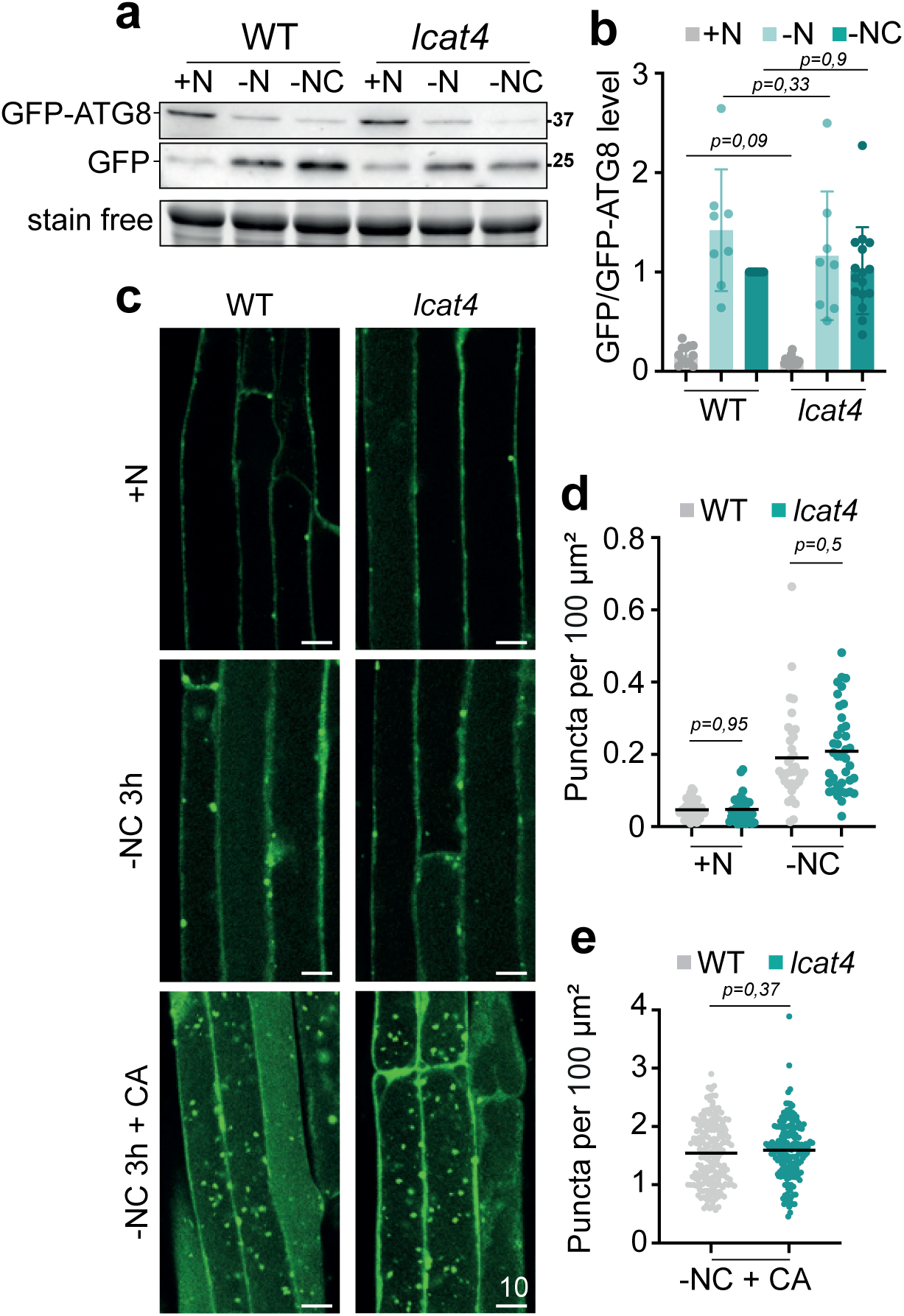
LCAT4 is dispensable for autophagy. **(a-b)** Autophagy flux is unaffected in the *lcat4* KO mutant compared to WT plants**. (a)** Detection of GFP-ATG8A degradation and release of free GFP of 7-day-old WT or *lcat4* plants expressing GFP-ATG8A. Seedling were transferred to either nutrient rich liquid medium (+N), nutrient starved liquid medium depleted in nitrogen (-N) or nitrogen and carbon (-NC) during 6h. Total proteins were extracted and analyzed by immunoblot with anti-GFP antibodies. Stain free images were used as loading control. **(b)** Quantification of the ratio of GFP/GFP-ATG8A in (a) relative to that of WT in -NC condition which was set to 1 in each experiment. Results are presented as the average, SD and values of all distinct replicates. In +N and -NC, n=12 (WT), n=16 (*lcat4*); in -N, n=8 (b). **(c-d)** The number of ATG8A-labeled compartments is not affected in the *lcat4* mutant compared to the WT**. (c)** Representative confocal images of WT or *lcat4* plants expressing GFP-ATG8A. Plants were imaged after 3 hours in rich condition (+N), deprived of nutrients for 3 hours (-NC 3h) or after 3h of -NC followed by 2 additional hours of -NC supplemented with concanamycin A (1 μM; -NC+CA). Scale bar, 10 µm. **(d)** Quantification of GFP-ATG8A puncta in conditions +N and -NC of (c). Results are presented as the number of puncta per 100 μm² of root area and show the average and individual values, from left to right, n=40 independent images over 10 roots; n=36 independent images over 9 roots; n=35 independent images over 9 roots and n=39 independent images over 10 roots. **(e)** Quantification of puncta labelled by GFP-ATG8A inside the vacuole in the condition -NC+CA of (c). Results are presented as the number of puncta per 100 μm^2^ of root area and show the average and individual values. For WT, n=178 cells and for *lcat4*, n=173 cells over 10 replicates. Statistical differences were assessed using two-tailed unpaired t-test (b, d, e).

### LCAT3, a potential redundant phospholipase for autophagy

The absence of a significant phenotype in the *lcat4* mutant prompted us to address the potential redundancy of additional phospholipases in the process of autophagic body degradation. LCAT4 belongs to a small multigenic family comprising four members: LCAT1, LCAT2 and LCAT3 (**Fig. S7a**). LCAT3 shares 53% identity with LCAT4 and was previously characterized as an acidic phospholipase A_1_ (**Fig. S7a, S8**; Noiriel et al., 2004). LCAT1 shares 23% identity with LCAT4, is a transmembrane protein previously found in a vacuolar proteome (**Fig. S7a**; Carter et al., 2004) and is predicted to have a hydrolase activity although it remains uncharacterized at this time. In contrast, LCAT2, also known as PSAT1, shares 20% identity with LCAT4 and has been previously demonstrated to have an acyltransferase activity involved in the formation of steryl esters (**Fig. S7a**; Shimada et al., 2021). Based on this data, we postulated that LCAT2 was unlikely to have a redundant function with LCAT4 and focused our research on LCAT3 and LCAT1.

Under stress-inducing conditions or during plant senescence, an increase in the expression of autophagy-related genes support autophagic induction and activity, notably improving stress tolerance (Agbemafle et al., 2023). To investigate whether LCAT3 and LCAT1 can participate in the autophagy pathway, we first analyzed their expression profiles in response to autophagy induction. First, the relative abundance of *LCAT1* and *LCAT3* transcripts was measured and compared to that of *LCAT4* and *ATG8A* during leaf development using RT-qPCR. Similar to *ATG8A*, *LCAT3* and *LCAT4* were found up-regulated during plant ageing; conversely, the transcripts of *LCAT1* decreased under the same conditions (**Fig. 4a**). In addition, we measured the expression level of *LCATs* genes in 7-day-old seedlings after 6 hours of nitrogen and carbon starvation. Similar to what we measured during senescence, we recorded an increase in the transcripts of *LCAT4* and *LCAT3*, and decreased abundance of *LCAT1* in these conditions (**Fig. S7b**). Based on the repression of *LCAT1* in autophagy-inducing conditions, we did not consider this gene further in this study. In contrast, the increased expression of *LCAT3,* alongside *LCAT4,* in conditions where autophagy is induced, either upon nutrient deficiency or senescence, supports its potential involvement in the autophagy pathway.

**Figure 4.**
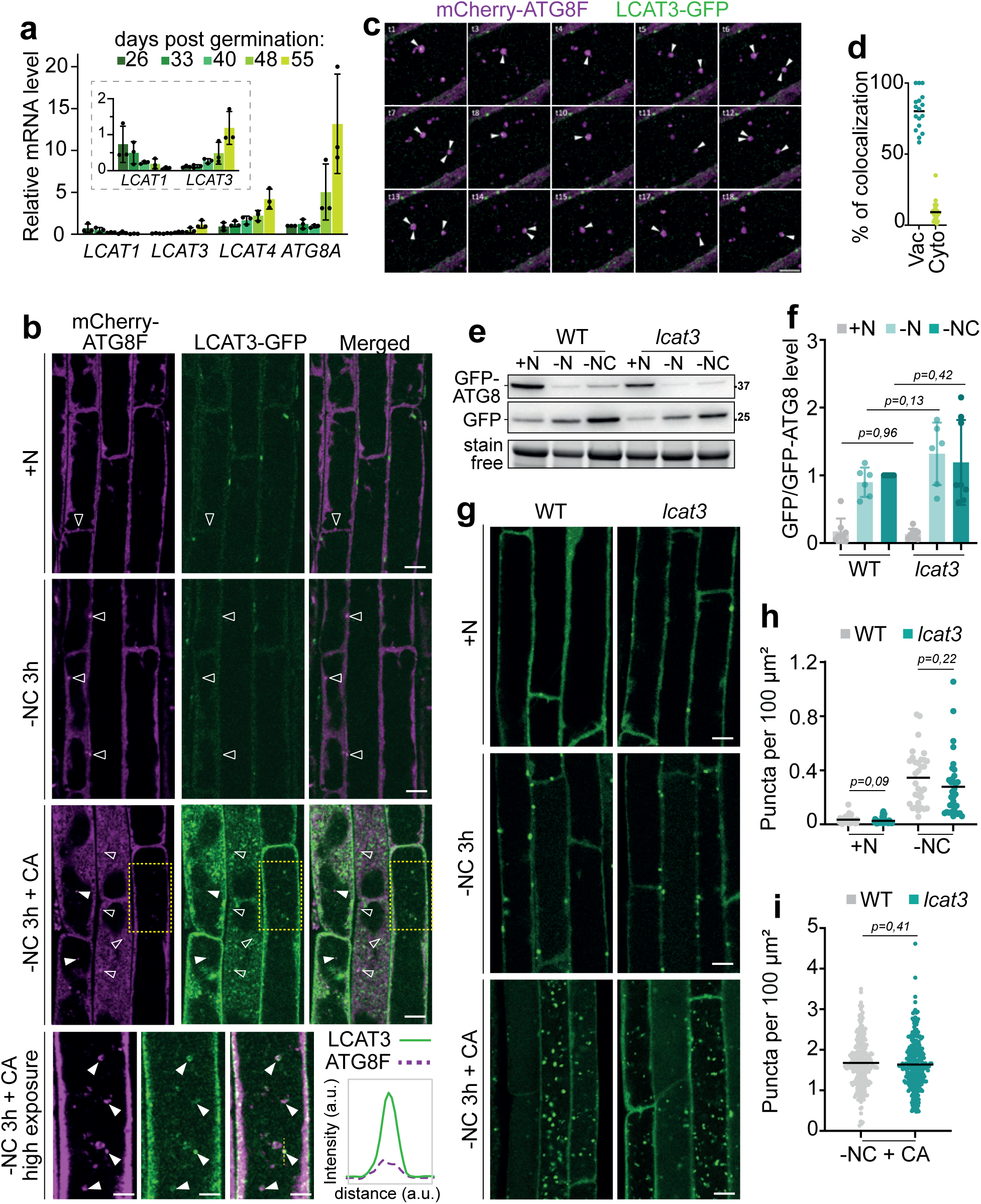
LCAT3 localizes on autophagic bodies in the vacuolar lumen but knocking out *LCAT3* has no effect on autophagy activity. **(a)** The expression of *LCAT3* increases during plant ageing, similar to that of *LCAT4* and *ATG8A*. Abundance of *LCAT1, LCAT3, LCAT4* and *ATG8A* transcripts in leaves of 26-, 33-, 40-, 48-and 55 day-old plants relative to *ATG8A* transcripts at day 26 which was set to 1 in each experiment. Histograms of *LCAT1* and *LCAT3* were enlarged in the insert to allow data visualization. Results show average +/-SD and all values of n=3 biological replicates. **(b-d)** The weak signal of LCAT3-GFP observed in +N or - NC conditions increases upon treatment with concanamycin A and shows co-localization with ATG8F on autophagic bodies in the vacuole. **(b)** Representative confocal images of roots of plants co-expressing LCAT3-GFP and mCherry-ATG8F. Plants were imaged after 3 hours in rich condition (+N), 3 hours of nutrient starvation (-NC 3h), or in -NC supplemented with concanamycin A (-NC 3h+CA, 1 μM). Empty arrowheads indicate autophagosomes in the cytosol and show the mostly distinct pattern of localization between the two proteins in these compartments. Full arrowheads point to autophagic bodies in the vacuole and show overlapping signals of ATG8F and LCAT3. Images of each condition are presented with the same settings showing the increased signal intensity of LCAT3-GFP in -NC+CA conditions compared to +N or -NC. For +N, -NC and -NC+CA, scale bar, 10 µm. Yellow dotted rectangles of the -NC+CA condition are enlarged on the lowest panel and the ATG8F signal was highly exposed to discern autophagic bodies in the vacuole; scale bar is 5 µm. Signal intensity profiles along the dotted lines are plotted in the boxed region on the right and show co-localization between ATG8F and LCAT3 puncta. **(c)** Representative time-lapse images of root cells imaged in -NC+CA conditions as in (b) showing constant association of LCAT3 puncta on ATG8F-labeled autophagic bodies in the vacuole over time. Time (in seconds) is indicated in the upper left corner; full arrows point to LCAT3 punctate signal. Sclae bar, 5 µm**. (d)** Quantification of colocalization events in (b). Results are presented as percentage of LCAT3-GFP puncta colocalizing with mCherry-ATG8F in the vacuole (Vac) or in the cytosol (Cyto). N=24 independent images taken over 12 roots (Vac) or 11 roots (Cyto) in 4 to 5 independent experiments. **(e-i)** Autophagy flux, autophagosome formation and autophagic body turnover are not affected in the *lcat3* KO mutant compared to WT plants. **(e)** Representative image of GFP-ATG8A processing assay performed as in Fig.3a in WT or *lcat3* plants expressing GFP-ATG8A. **(f)** Quantification of the ratio of GFP/GFP-ATG8A of (e) relative to that of WT in (-NC) condition which was set to 1 in each experiment. Results are presented as the average ± SD with values of each independent biological experiments; +N or -NC, n=8; -N, n=6. **(g)** Representative confocal images of WT or *lcat3* plants expressing GFP-ATG8A. Plants were imaged as in Fig. 3c. Scale bar, 10 μm. **(h)** Quantification of GFP-ATG8A puncta in conditions +N or -NC of (g). Results are presented as the number of puncta per 100 μm^2^ of root area and show the average and individual values of n=number of images in N=number of independent plants in two independent experiments. From left to right: n=40, N=12; n=40, N=10; n=32, N=9; n=33, N=9. **(i)** Quantification of GFP-ATG8A puncta in the vacuole in -NC 3h+CA of (g). Results are presented as the number of puncta per 100 μm^2^ of root area and show the average and individual values of n=253 cells over 13 independent plants in 2 biological replicates (WT) and n=272 cells over 11 independent plants in 2 biological replicates for *lcat3*. Statistical differences were assessed using two-tailed unpaired t-test (f, h, i).

To test the whether LCAT3 participates in the autophagy pathway, we analyze the subcellular distribution of the protein and autophagy compartments by generating stable *Arabidopsis* lines co-expressing LCAT3-GFP and mCherry-ATG8F. Imaging the roots of 7-day-old seedlings, under rich or -NC conditions using confocal microscopy, we observed that the LCAT3-GFP signal was very weak, even when the transgene was expressed under the control of the constitutive pUBQ10 promoter (**Fig. 4b**). In these conditions, LCAT3-GFP was mostly found in the cytosol or on cytosolic puncta that do not co-localize with mCherry-ATG8F. This result indicates that LCAT3 is not present on autophagosomes, in contrast to LCAT4 which was found to be recruited to early autophagy compartments (**Fig. 1**, **Fig. S1**). This difference in localization between the two proteins may be specified by their diverging C-terminal parts: although they show high sequence identities overall, LCAT3 is shorter and lacks the C-terminal tail of LCAT4 where the predicted ATG8 interacting domain is located (**Fig. S8a-b**). Accordingly, AlphaFold3 models did not predict an interaction between LCAT3 and ATG8 (**Fig. S8a-d**) which is consistent with results in **Fig. 1a** where LCAT3, in contrast to LCAT4, was not found enriched in ATG8-immunoisolated compartments.

In contrast to -NC conditions, where LCAT3 did not co-localize with ATG8, when plants were additionally treated with concanamycin A (which enables the visualization of GFP signal in the vacuole, otherwise quenched by the acidity of its lumen) we observed LCAT3 inside the vacuole, on bright foci associated with ATG8-labeled compartments (**Fig. 4b-d**). This result indicates that LCAT3 traffics to the vacuolar lumen and localizes on autophagic bodies. The absence of LCAT3 on autophagosomes coupled with the presence of LCAT3 in the vacuole suggests that, in contrast to LCAT4, LCAT3 traffics to the vacuole through the canonical pathway for vacuolar resident proteins, *i.e.,* debuting in the ER, passing first through the Golgi, then the Trans-Golgi-Network (TGN) and finally the MVB prior to reaching the vacuole lumen (Aniento et al., 2022). This hypothesis is supported by the large increase in the overall level of LCAT3-GFP in +CA conditions, with the protein observed on numerous small mobile puncta in the cytosol that do not, or rarely, colocalize with ATG8 (**Fig. 4b,d**). TGN maturation and MVB trafficking is partially blocked by addition of concanamycin A (Dettmer et al., 2006), therefore the accumulation of cytosolic LCAT3 puncta in these conditions could reflect its transient trafficking through these compartments in untreated conditions. Despite this trafficking route remaining to be characterized, our results clearly demonstrate that a portion of LCAT3 localizes in the vacuole lumen, likely independently of autophagy, and can recognize and associate with autophagic bodies.

### LCAT4 and LCAT3 control autophagy degradation in the vacuole

The homology between LCAT4 and LCAT3, as well as their localization on autophagic bodies, suggest that these two phospholipases might have redundant or synergistic roles in autophagy. To test this hypothesis, we collected a *lcat3* T-DNA KO mutant, generated a conditional *LCAT3* KD mutant and constructed a double *lcat3 lcat4* KO mutant by crossing *lcat3* and *lcat4* T-DNA insertion lines (**Fig. S5a-c).** To measure autophagy, the GFP-ATG8A construct was introgressed in the aforementioned lines which were used to perform the GFP-ATG8 degradation assay and to quantify the number of autophagy compartments *in vivo*. In the *lcat3* KO line, we did not observe significant defects in autophagic flux (**Fig. 4e-f**) or in the number of GFP-ATG8A-labeled autophagosomes or autophagic bodies (**Fig. 4g-i**). In contrast, when the expression of *LCAT3* was reduced for a few hours prior to autophagy induction, in the conditional *amiRNA:LCAT3* line, we detected a slight decrease in autophagic flux in -N conditions (**Fig. S9**). These results suggest that LCAT3 has a function in the autophagy pathway and that its complete absence in the *lcat3* KO line is likely complemented by a redundant activity.

In contrast to single *lcat3* or *lcat4* mutants, analyses of the autophagy flux using the GFP-ATG8 processing assay in the double *lcat3 lcat4* KO line (hereafter *lcat3,4*), showed a decrease in the GFP/GFP-ATG8 ratio, approximately 50% lower than in WT, under either nitrogen- or combined nitrogen and carbon-depleted conditions (**Fig. 5a-b**). These results show that knocking out both *LCAT3* and *LCAT4* slows down autophagy activity and suggest that LCAT3 and LCAT4 have redundant and/or synergistic effects in the autophagy pathway. To determine the stage of the pathway at which LCAT3 and LCAT4 participate during autophagy, we analyzed the number and distribution of autophagic structures in the *lcat3*,*4* double mutant. We assessed autophagosome formation, by counting the number of ATG8A puncta in the cytosol in rich conditions (+NC) or in autophagy-inducing conditions (-NC, 3h). As seen in (**Fig. 5c,e)**, these analyses showed no significant differences in the *lcat3,4* double mutant compared to WT plants. This shows that the autophagy defects observed in the absence of LCAT3 and LCAT4 are not caused by a reduction in autophagosome formation or by an inhibition of autophagosome to vacuole trafficking. Strikingly, these analyses revealed a major difference between *lcat3,4* and WT plants in the vacuole: in -NC conditions, we observed an accumulation of GFP-ATG8A puncta inside the vacuolar lumen of the *lcat3,4* double mutant which are barely ever observed in cells of WT plants (see -NC conditions in **Fig. 5c-f**). This phenotype was recapitulated in an alternative autophagy-inducing condition, *i.e.*, upon treatment with the TOR inhibitor AZD-8055 (AZD, **Fig. S10**) demonstrating the robustness of this result. Additionally, when concanamycin A was further added after 3 hours of prior autophagy induction, the number of GFP-ATG8A puncta in the vacuole increased in both genotypes with a significant greater abundance in the double *lcat3,4* mutant compared to WT (see -NC +CA conditions in **Fig. 5c,e** and similar phenotype upon AZD+CA treatment in **Fig. S10**). These results, in contrast to the deletion of either *LCAT4* or *LCAT3* which showed no impact on autophagy compartments **(Fig. 3c,e** and **Fig. 4g,i**, respectively), reveal that the *lcat3,4* mutant accumulates autophagic bodies within the vacuole, indicating that their proper degradation requires LCAT3 and LCAT4 and suggesting partial redundant activities of the proteins. Further, the concomitant accumulation of autophagic bodies and reduction of autophagy flux in the *lcat3,4* mutant demonstrate the importance of autophagic body disruption for the completion of autophagy. Nevertheless, the 50% remaining autophagy activity detected in the *lcat3,4* mutant (**Fig. 5a,b**) suggests that additional proteins, yet to be identified, are involved in this process. In fact, compared to WT plants, the *lcat3,4* mutant showed similar germination, root length, vegetative growth or onset of senescence, unlike *atg* mutants where autophagy is completely abolished (**Fig. S11**). These results show that the absence of the two proteins, and the corresponding 50% reduction in autophagy flux has no major effect on plant development in rich conditions. In contrast, we observed an intermediate phenotype between that of *atg5* and that of WT when *lcat3,4* seedlings were placed in -NC for 10 days (**Fig. 5g,h**). While none of the *atg5* mutants were able to cope with prolonged starvation, most WT seedlings were found to be mostly green when the *lcat3,4* mutant displayed a mild yet significant senescing phenotype. Together, these analyses show that LCAT3 and LCAT4 are core components of the autophagy machinery, and participate, or control, the hydrolysis of autophagic bodies at the antepenultimate step of the pathway.

**Figure 5.**
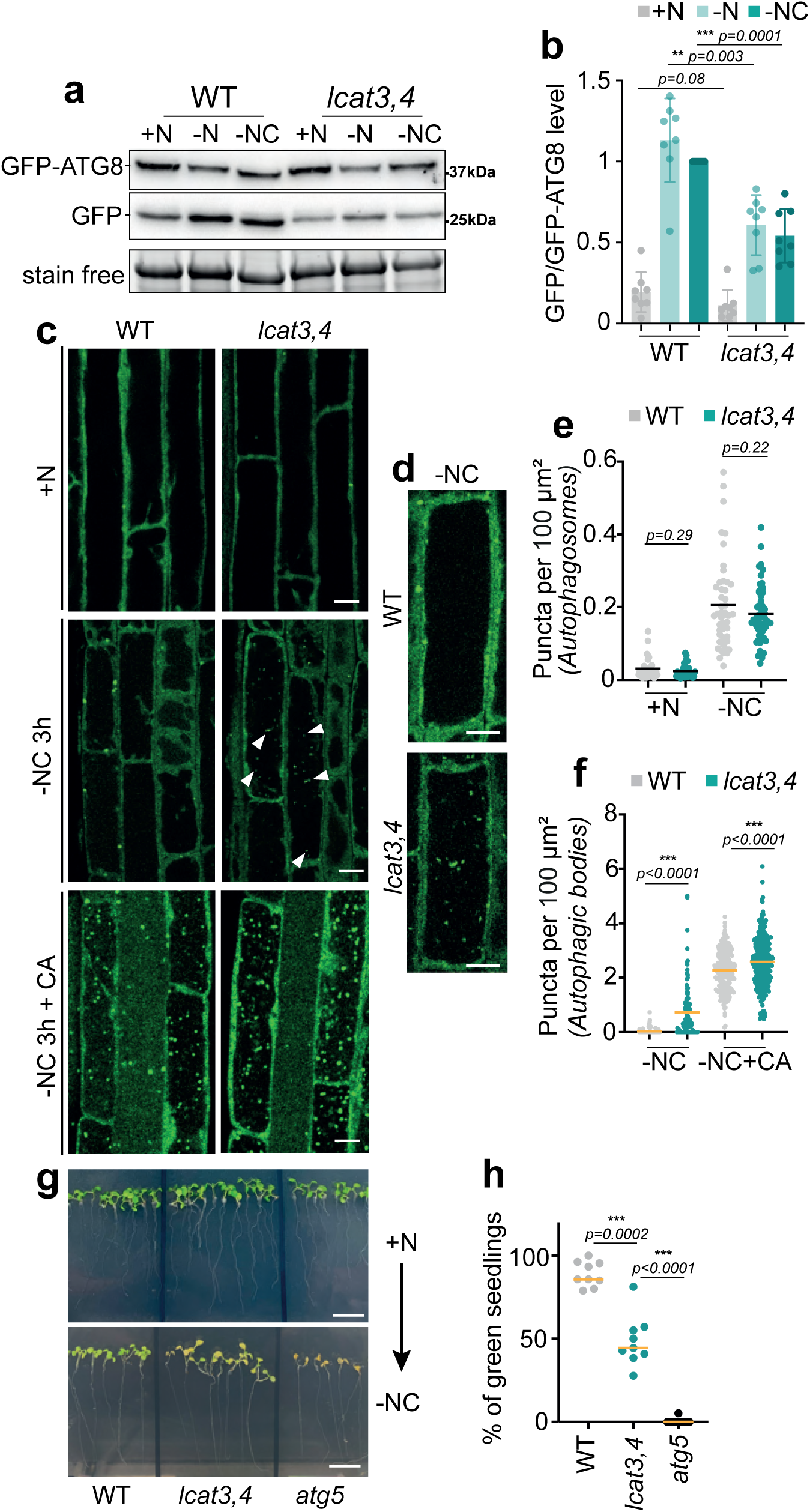
Knocking out both *LCAT4* and *LCAT3* causes defects in autophagy activity and a concomitant accumulation of autophagic bodies. **(a-b)** Autophagy flux is significantly decreased in the *lcat3,4* double mutant compared to WT plants. **(a)** Representative image of GFP-ATG8A processing assay performed as in Fig.3a in WT or *lcat3,4* double knock out plants expressing GFP-ATG8A. **(b)** Quantification of the ratio of GFP/GFP-ATG8A of (a) relative to that of WT in (-NC) condition which was set to 1 in each experiment. Results are presented as the average ± SD with values of all replicates, n=8 independent biological experiments for each condition. **(c-f)** Autophagic bodies accumulate in the *lcat3,4* mutant compared to WT plants. **(c)** Representative confocal images of WT or *lcat3,4* plants expressing GFP-ATG8A. Plants were imaged as in Fig. 3c. Scale bar, 10 μm. Arrowheads point to ATG8A-labbeled autophagic bodies in the vacuole of *lcat3,4* in -NC conditions. **(d)** Representative image of a root cell of *lcat3,4* in -NC condition showing an accumulation of GFP-ATG8A puncta in the vacuole compared to the empty vacuole of a WT cell in the same condition. **(e)** Quantification of GFP-ATG8A puncta in +N or -NC of (c). Results are presented as the number of puncta per 100 μm^2^ of root area and show the average and individual values of n=number of images in N=number of independent plants in B=number of independent biological experiments. From left to right: n=35, N=12, B=2; n=32, N=11, B=2; n=47, N=16, B=3 and n=58, N=18, B=3. **(f)** Quantification of autophagic bodies (*i.e.*, ATG8A puncta in the vacuole) in -NC or -NC+CA conditions of (c). Results are presented as the number of autophagic bodies per 100 μm^2^ of root area and show the average and individual values of n=number of cells in N=number of independent plants in B=number of independent biological experiments. From left to right: n=121, N=10, B=3; n=115, N=11, B=3; n=233, N=9, B=2 and n=251, N=10, B=2. **(g-h)** Compared to WT plants, the *lcat3,4* double knock out mutant is less tolerant to prolonged nutrient starvation. Seedlings were grown on MS + Sucrose for 7 days (g, +N, upper panel) before being transferred to -NC condition for 10 days (g, -NC, lower panel). **(g)** Representative images; scale bar, 1 cm. **(h)** Qualitative quantification of (g). Results are expressed as percentage of mostly green seedlings compared to total seedlings for each genotype. A total of 161 (WT), 143 (*lcat3,4*) and 142 (*atg5*) seedlings were analyzed in 9 independent replicates. Results are presented as the average and individual values for each independent replicate. Statistical differences were assessed using the Mann Whitney test (b, h) or two-tailed unpaired t-test (e, f).

### LCAT3 and LCAT4 act synergistically to disrupt the membrane of autophagic bodies

Once the autophagic bodies are delivered inside the vacuolar lumen, the disruption of their membrane is a *sine qua non* condition for autophagy progression: it enables the release of the cargo for subsequent degradation and recycling, thereby completing autophagy activity. Further, this process must be tightly regulated by specific lipid-hydrolytic enzymes, able to specifically recognize and target the membrane of autophagic bodies without affecting that of the vacuole, the tonoplast, to keep cellular homeostasis. Based on the localization of LCAT3 and LCAT4 (**Fig.4, 1**), on the nature of their enzymatic activity as phospholipases (Noiriel et al., 2004; Chen et al., 2012) and on their impact on the autophagy pathway (**Fig. 5**) we hypothesized that they participate in the hydrolysis of the membrane of autophagic bodies. To test this potential function, we set up a system in which LCAT3 and LCAT4 were expressed in yeast cells that are genetically impaired for the degradation of autophagic bodies. In the budding yeast *S. cerevisiae*, a single protein called Atg15, a phospholipase B, is required for the breakdown of autophagic bodies (Kagohashi et al., 2023; Watanabe et al., 2023). Although there is no sequence homologue for Atg15 in plants (Bassham et al., 2006), we proposed to challenge our idea that LCAT3 and LCAT4 could be functional homologues of this phospholipase in autophagy by testing whether their expression can complement the absence of Atg15 in yeast. To monitor the disruption of autophagic bodies in LCAT3- or LCAT4-expressing *atg15Δ* cells, we first followed the level of the mature form of the vacuolar-resident aminopeptidase 1, Ape1. Ape1 uses a selective type of autophagy, called the Cvt pathway (Cytoplasm-to-Vacuole Targeting), to traffic to the vacuole; inside the vacuolar lumen the prApe1 propeptide is cleaved by vacuolar proteases thus converting Ape1 into a mature and active form (mApe1). In the absence of Atg15, prApe1 reaches the vacuolar lumen but remains entrapped inside autophagic bodies, preventing its maturation (Kagohashi et al., 2023 and **Fig. 6a-b**, compare the empty plasmid condition (e.p.) to WT cells). Expressing either *LCAT3* or *LCAT4* in *atg15Δ* cells did not result in the formation of mApe1, indicating that, at steady state, the proteins cannot rescue the *atg15Δ* phenotype, *i.e.,* cannot disrupt autophagic bodies (**Fig. 6a-b**). In contrast, when LCAT3 was fused to the signal sequence of carboxypeptidase Y (CPY) which addresses protein inside the vacuolar lumen (Kagohashi et al., 2023 and **Fig. S12**), we detected the mature form of mApe1 (mApe1) indicating that a portion of autophagic bodies were hydrolyzed (**Fig. 6a,b**). Previous work on LCAT3 identified its catalytic triad, consisting of S177, D384, and H409 and showed that replacing S177 to an alanine prevented the phospholipase activity of the protein (Noiriel et al., 2004). Compared to CPY-LCAT3^WT^, *atg15Δ* cells expressing CPY-LCAT3^S177A^ failed to accumulate mApe1 indicating that the phospholipase activity of LCAT3 is required for its capacity to breakdown autophagic bodies (**Fig. 6c**). From these results we concluded that, when expressed in the vacuole, LCAT3 can recognize and hydrolyze the membrane of autophagic bodies in yeast. In contrast, when expressing LCAT4, even when it was fused to the CPY vacuolar-addressing peptide (**Fig. S12**), we did not detect mApe1 (**Fig. 6a,b**), showing that LCAT4 alone cannot complement the *atg15Δ* phenotype and disrupt the membrane of autophagic bodies.

**Figure 6.**
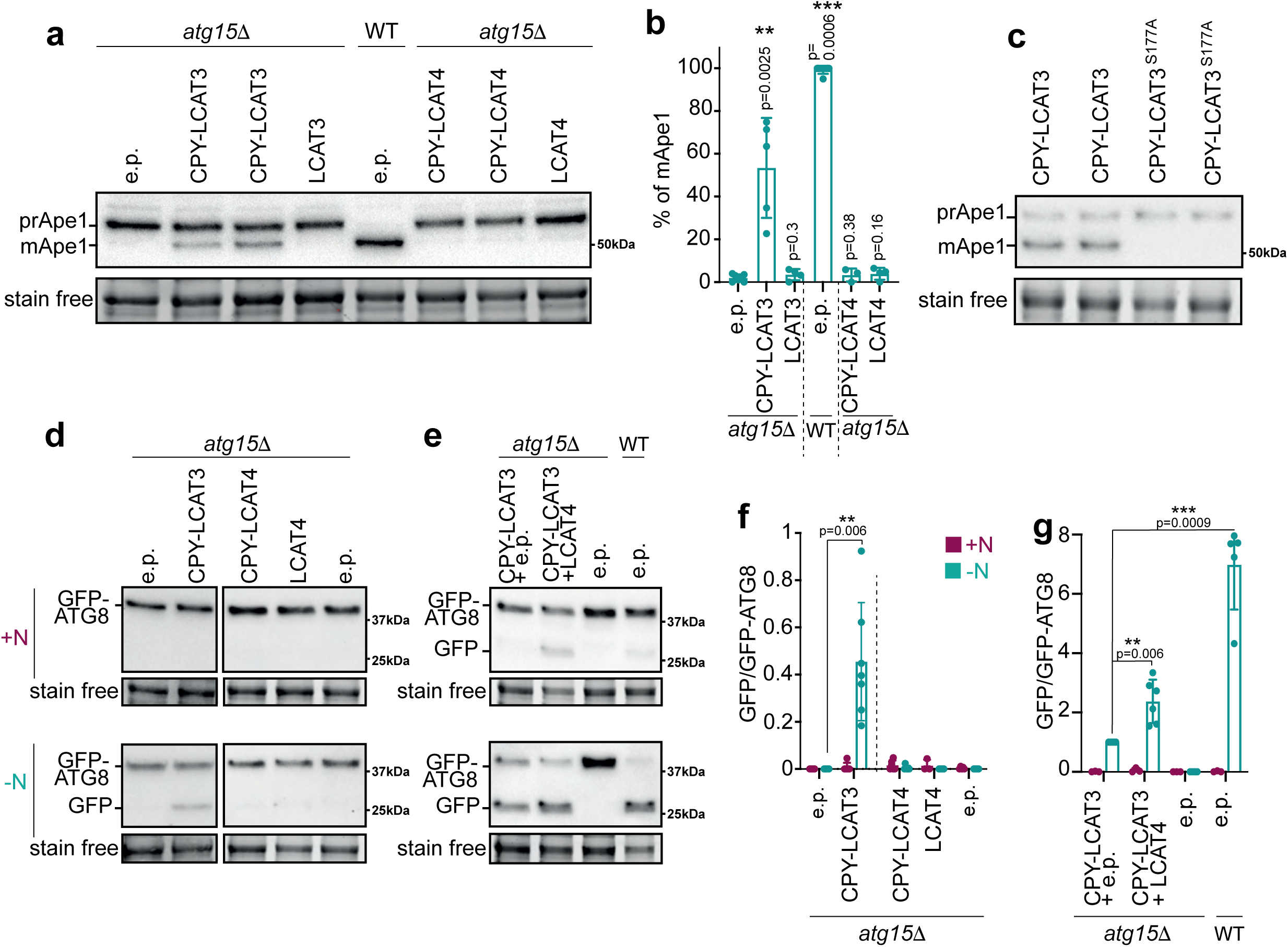
LCAT3 and LCAT4 can disrupt the membrane of autophagic bodies. **(a-c)** The phospholipase activity of LCAT3 can partially complements the phenotype of *atg15Δ* cells. **(a)** Representative image of the immunoblot analyses of Ape1 in WT or *atg15Δ* cells transformed with the empty vector (e.p.) or expressing CPY-LCAT3 (2 independent replicates), LCAT3, CPY-LCAT4 (2 independent replicates) or LCAT4. Cells were grown until mid-log phase in SMD-TRP+GAL conditions prior to protein extraction and immunoblot analyses. **(b)** Quantification of the percentage of mApe1 compared to prApe1 from (a). Results are presented as average +/- SD with individual values of each independent replicate, from left to right: n=7, n=5, n=4, n=7, n=3, n=4. Statistical differences between *atg15Δ* + e.p. and other conditions were assessed using Mann Whitney tests. **(c)** Immunoblot analyses of Ape1 in *atg15Δ* cells transformed with either CPY-LCAT3 or CPY-LCAT3^S177A^. Results of 2 independent replicates are shown. **(d-g)** Expressing CPY-LCAT3 partially restores autophagy degradation in *atg15Δ* cells (d,f) and autophagy flux is further enhanced when CPY-LCAT3 and LCAT4 are co-expressed (e,g). **(d,e)** Representative images of the immunoblot analyses of GFP-AtATG8A processing assays in *atg15Δ* yeast transformed with the empty vector (e.p.), CPY-LCAT3, CPY-LCAT4 or LCAT4 (d) or co-transformed with CPY-LCAT3+e.p or CPY-LCAT3+LCAT4 compared to WT cells transformed with the corresponding empty plasmids (e). Yeast cells were grown in SMD+GAL selective media (-TRP for e.p. and CPY-LCAT3, d, left panel; -URA for CPY-LCAT4, LCAT4 and e.p., d, right panel; -TRP-URA for CPY-LCAT3+e.p. and CPY-LCAT3+LCAT4, e) until mid-log phase (+N) prior to be transferred to nutrient starvation (-N, SD-N+GAL). Stain free images were used as loading control. **(f)** Quantification of the ratio of GFP/GFP-ATG8A of (d). Results are presented as the average ± SD with values of each independent replicates, for each histogram, from left to right: n=4, 4, 7, 7, 3, 4, 6, 6, 5, 6. Statistical differences between *atg15Δ* + e.p. and other conditions were assessed using Mann Whitney tests, significant p values are indicated. **(g)** Quantification of the ratio of GFP/GFP-ATG8A of (e) relative to that of CPY-LCAT3 in (-N) condition which was set to 1 in each experiment. Results are presented as the average ± SD with values of each independent replicates, for each histogram, from left to right: n=3, 6, 3, 6, 3, 5, 3, 5. Statistical differences between CPY-LCAT3 + e.p. in -N and other conditions were assessed using one sample t.test, significant p values are indicated.

From our results in **Fig.1-2**, we hypothesized that LCAT4 is recruited to the phagophore membrane by its interaction with ATG8 and traffics inside the autophagosomes/autophagic bodies to reach the vacuolar lumen. In contrast, we found that LCAT3 traffics to the vacuole independently of autophagy (**Fig. 4b,c**) and can recognize the outer leaflet of the membrane of autophagic bodies either in *Arabidopsis* (**Fig. 4b,c**) or in yeast cells (**Fig. 6a,b**). Based on this data, we postulated that LCAT3 and LCAT4 act synergistically, with LCAT3 starting from the outer leaflet, providing initial membrane disruption to activate LCAT4 localized in the inner leaflet of the vesicular membrane. To test this hypothesis, we swapped the endogenous Atg8 from *S. cerevisiae* to GFP-ATG8A from *Arabidopsis* at the Atg8 locus and tested the impact of iterative expression of CPY-LCAT3 and LCAT4 on GFP-ATG8A processing, as a proxy for autophagic body disruption and autophagy degradation. Our results recapitulated the observation from the analyses of Ape1 in **Fig.6a,b** by showing that CPY-LCAT3 promotes the degradation of GFP-ATG8A resulting in the accumulation of free GFP upon autophagy induction by nitrogen starvation (**Fig. 6d,f**; left panel) while expressing either LCAT4 or CPY-LCAT4 did not result in GFP-ATG8A processing (**Fig. 6d,f**; right panel). However, when we mimicked plant trafficking/localization scenarios, by co-expressing GFP-ATG8A and LCAT4 (in the cytosol) together with CPY-LCAT3 (in the vacuole) in *atg15Δ* cells, we observed a significant increase in autophagy activity compared to the expression of CPY-LCAT3 alone (**Fig. 6e,g**). This result indicates that both LCAT3 and LCAT4 can hydrolyze the membrane of autophagic bodies in yeast cells. It further supports the hypothesis that the activity of LCAT4 requires that of LCAT3 and suggests that both enzyme act synergistically to instruct the disruption of autophagic bodies as a prerequisite for the completion of autophagy degradation.

## Discussion

### LCAT3 and LCAT4 catalyze the antepenultimate step of the autophagy pathway

The degradation and recycling of proteins, organelles and pathogens through the autophagy pathway play critical roles in plant development and plant tolerance to environmental stresses (Gross et al., 2025). Yet, while our understanding of the formation, cargo packing and trafficking of autophagy vesicles has greatly increased (Zhuang et al., 2024), what occurs inside the vacuole during the actual degradation and recycling steps of the autophagy pathway remain uncharacterized in plants and poorly described across models.

Here we uncover the role of two phospholipases, LCAT3 and LCAT4, in the disruption of autophagic bodies in the vacuole (**Fig. 5, 6**). We show that preventing this process leads to a large decrease in autophagy flux (**Fig. 5**) supporting the critical relevance of this membrane remodeling event in the completion of the autophagy pathway. Therefore, we propose that both LCAT3 and LCAT4 are part of the core machinery of autophagy. From our results we further draw a model according to which plants have evolved a multi-component mechanism to control, spatially specify, and efficiently instruct the disruption of autophagic bodies. Our results show that LCAT4 is mostly found in the cytosol at steady state (**Fig. 1**), *i.e.*, in conditions of neutral pH at which the protein was found inactive *in vitro* (Chen et al., 2012). LCAT4 directly interacts with ATG8, is recruited to the phagophore membrane and remains associated with autophagosomes within which it traffics to the acidic vacuolar lumen upon autophagy induction (**Fig. 1, 2**). This places LCAT4 in the inner leaflet of the membrane of autophagic bodies. In contrast, LCAT3 does not localize to autophagosomes yet traffics to the vacuolar lumen where it recognizes and associates with autophagic bodies (**Fig. 4**), which indicates that LCAT3 associates with the outer leaflet of the membrane of autophagic bodies. When targeted to the vacuolar lumen in yeast cells, LCAT3 can recognize and break down autophagic bodies (**Fig. 6**), which is consistent with previous work showing the requirement of an acidic environment for the phospholipase activity of LCAT3 (Noiriel et al., 2004). However, *lcat3* KO/KD mutants show little to no difference in the rate of autophagy degradation in *Arabidopsis* indicating that the function of LCAT3 is complemented by additional or redundant activities. In contrast to single *lcat3* or *lcat4* mutants, the double *lcat3,4* knock out shows a large reduction in autophagy flux and a concomitant accumulation of autophagic bodies in the vacuole (**Fig. 5**). We thus propose a model in which a synergy between LCAT3 and LCAT4 is required to efficiently promote the degradation of autophagic vesicles. In that model, LCAT4 is transported and trapped inside autophagic bodies and requires LCAT3 to perform the initial hydrolyzing step, destabilizing these compartments and thus enabling LCAT4 to become active at acidic pH. A combination of both activities increases the disruption of autophagic bodies when LCAT3 and LCAT4 were expressed in yeast (**Fig. 6**), supporting the requirement and synergy of the proteins to efficiently process autophagic bodies in the vacuole. Finally, we note that plants lacking LCAT3 and LCAT4 are still able to perform autophagy, albeit at a lower rate (**Fig. 5**). This indicates that additional phospholipases act redundantly (or compensate LCAT3/LCAT4) in this pathway. Our study here opens the way to the understanding of the regulation of autophagic bodies turn-over and our current work is focusing on the identification and characterization of additional components of the antepenultimate step of autophagy.

### A two-component system to instruct the efficient degradation of autophagic bodies?

With the exception of Atg15, a phospholipase B which controls this step of the pathway in yeast cells, little is known about the processing of autophagic bodies membranes across organisms. While yeast and plant cells show similar features in their lytic compartments, with large and unique vacuoles compared to the small and numerous lysosomes in mammals, LCAT3 and LCAT4 do not resemble Atg15, which has no sequence homologs in *Arabidopsis*. In contrast, they show sequence homologies with lysosomal phospholipases from human (PLA_2_G15; phospholipase A_2_ group XV) or from *C. Elegans* (LPLA-2) which have been involved in the degradation of intralumenal lysosomal material (Shayman et al., 2011; Li et al., 2022). Nevertheless, the expression of LCAT3 in the vacuole of yeast cells can partially complement the absence of Atg15, suggesting functional homologies with the yeast protein. As mentioned above, recent work found that Atg15 has a phospholipase B activity, *i.e.*, hydrolyze both acyl chains from phospholipids (Watanabe et al., 2023; Kagohashi et al., 2023). In contrast, LCAT3 and LCAT4 were found to have phospholipase A activities, hydrolyzing the acyl chain in *sn-1* and *sn-2* positions of phospholipids, respectively (Noiriel et al., 2004; Chen et al., 2012). In that context, the combined action of LCAT3 and LCAT4 could mimic that of a phospholipase B and might be a prerequisite for the efficient hydrolysis of phospholipids in the membrane of autophagic bodies.

### A two-component system to instruct the tight spatiotemporal control of autophagic body degradation?

While the membrane of autophagic bodies needs to be broken off to ensure cargo delivery and degradation, this step must be tightly regulated in a spatiotemporal manner. In time, to efficiently process the large influx of autophagic bodies upon autophagy induction, and in space, to hydrolyze the membrane of autophagic bodies without affecting the homeostasis of the tonoplast which defects can lead to cell death (van Doorn et al., 2011). How this regulation is achieved remains an open and critical question that our work starts to address.

First, our study here shows that *LCAT4* and *LCAT3*, similarly to other *ATG* genes, are upregulated upon autophagy inducing conditions (**Fig. 4a**; Agbemafle et al., 2023). This suggests that the rate of autophagic bodies’ turn-over is under transcriptional control. Second, we propose that the disruption of autophagic bodies is spatially regulated by the trafficking of LCAT3 and LCAT4 to the vacuole and pH-dependent activity of these proteins. Previous *in vitro* characterization of the phospholipase activity of each protein showed that both are only active in acidic pH (Chen et al., 2012; Noiriel et al., 2004). This supports that LCAT3 and LCAT4 are inactive in the pH neutral cytosol even though LCAT4 is already associated with autophagy compartments. Once in the acidic sap of the vacuolar lumen, both proteins become active thereby spatially compartmentalizing the degradation of autophagic bodies. Other organelles of the cells (TGN, MVB) show an acidic pH, albeit more alkaline than that of the vacuole (Shen et al., 2013). Depending on the trafficking route of LCAT3, we cannot exclude that LCAT3 may already become active prior to reaching the vacuolar lumen, and may therefore initiate membrane hydrolysis during autophagosome maturation as amphisomes (upon autophagosome/MVB fusion; Zhao et al., 2022) or in the recently characterized VAPV (VPS41-Associated Phagic Vacuoles) in which partial degradation was suggested (Jiang et al., 2004). In yeast, Atg15 uses the MVB pathway to reach the vacuole through its transmembrane domain, which acts as a signal sequence (Watanabe et al., 2023; Kagohashi et al., 2023). Analyses of the amino-acid sequence of LCAT3 did not predict a specific signal peptide or a conserved vacuole sorting determinant. It is therefore difficult to speculate on the pathway used by LCAT3 to reach the vacuole. Yet, the accumulation of LCAT3 cytosolic puncta upon treatment with concanamycin could reflect its transient passing through the TGN and/or MVBs as TGN maturation and MVB trafficking is partially blocked in these conditions (Dettmer et al., 2006). Further, a proportion of autophagosomes fuses with MVBs prior to their delivery to the vacuole (Zhao et al., 2022) which could be the point of contact between LCAT3 and the autophagy compartments. Supporting this idea, our analyses of the subcellular localization of LCAT3 found transient proximity between the cytosolic pool of LCAT3-marked vesicles and autophagosomes close to the tonoplast (**Movie S1**). Future research will thus aim at elucidating the transport pathway of this protein to the vacuole.

Third, we propose that the specific localization of LCAT3 and/or LCAT4 at the membrane of autophagic bodies spatially determines the site of their activity, thereby preventing tonoplast hydrolysis. While the molecular determinants for such localization remain unknown for LCAT3, the protein was found associating with microsomes when expressed in yeast cells (Noiriel et al., 2004) and we showed here that LCAT3 can hydrolyze autophagic bodies when expressed in the yeast vacuole, demonstrating its ability to associate with the membrane of autophagic bodies even in a heterologous system (**Fig. 6**). These results suggest that LCAT3 can directly bind lipids/membranes specifically at autophagic bodies and we recognize at least three factors that could contribute to its distinct recruitment at autophagic vesicles inside the vacuole rather than at the tonoplast. (i) The singularity of their respective lipid composition, with phospholipids representing more than 90% of the total lipids of autophagy compartments (in *Arabidopsis*, based on our recent unpublished findings) when it makes up only 51% of the vacuolar membrane (in seedlings of Mung Bean, Yoshida and Uemura, 1986; note that the comprehensive lipid composition of the *Arabidopsis* tonoplast is unknown). (ii) The specific protein composition of each of these compartments is a potential alternative or additional layer of protection for the tonoplast, which may specify the recruitment of LCAT3 at the autophagic bodies. (iii) The morphology, in particular the membrane curvature, of these compartments, is also highly dissimilar and could participate in phospholipases distinction. Vacuolar membranes are generally characterized by very low or negative curvature (particularly in the elongated zone of the root where vacuoles form large bodies), whereas autophagic bodies have a high membrane curvature. In fact, other ATG proteins can sense membrane features such as membrane curvature or lipid packing defects. For example, mammalian ATG14 or yeast Atg1 can recognize membrane curvature (Fan et al., 2011; Chan et al., 2009; Ragusa et al., 2012). Recent work demonstrated that Atg15 can bind autophagic bodies membrane but not the tonoplast, explaining why this phospholipase is prevented to digest the vacuolar membrane in yeast event though the molecular mechanisms for such membrane specificity remain unknown (Kagohashi et al., 2023). In that context, similarly to Atg15, LCAT3 may contain membrane/lipid binding sites specific to either the composition or the biophysical properties of the autophagic bodies that future research should investigate in detail.

In contrast to LCAT3, previous work showed that the phospholipase activity of LCAT4 is mostly associated with the soluble fraction when expressed in yeast in rich conditions (Chen et al., 2012) which suggests that LCAT4 itself is unable to bind membranes. Here, we found that LCAT4 directly interacts with ATG8 in *Arabidopsis*, and thus propose that this protein/protein interaction recruits LCAT4 to its target membrane, thereby specifying the site of action of this hydrolase at autophagic bodies. Further work, outside the current study, is in progress to address this hypothesis.

### Autophagy as a transport system for vacuolar enzymes in plants

In addition to identifying the actors of autophagic bodies hydrolysis, our work here also finds that LCAT4 uses autophagy to traffic from the cytosol to the vacuole upon nutrient depletion (**Fig. 2**). Based on this data, we propose an unsuspected conceptual leap in the field, showing that membrane hydrolysis is decided as early as its formation in the cytoplasm, during autophagosome biogenesis. This trafficking activity of autophagy puts the enzyme at the right place, *i.e.*, at the membrane of autophagic bodies; at the right time, being transported inside the very vesicle that needs to be hydrolyzed; and at the same rate as their delivery inside the vacuole, to promote the efficient degradation of autophagic bodies. This data thus points to the somewhat unsuspected role of autophagy as a transport route in plants, prepping the vacuole to deal with the massive influx of autophagic bodies by packing up the hydrolytic enzyme(s) that will later be activated inside the vacuole and needed for its degradation and potentially that of its cargo.

In that respect, the pathway that we identified here resembles that of the cytoplasm-to-vacuole targeting (Cvt) pathway, a selective type autophagy in *Saccharomyces cerevisiae*, which transports hydrolases to the vacuole and notably the aminopeptidase Ape1 (Yamasaki and Noda, 2017). A similar transport route has never been identified in plants, where autophagy has solely been described as a degradation pathway (Gross et al., 2025). Yet, a recent study described that, when overexpressed in *N. benthamiana*, the cysteine proteases VPEs from potato traffic to the vacuole using autophagy under carbon starvation (Teper-Bamnolker et al., 2021). This example diverges from the Cvt pathway, because even though autophagy is used as a mean of transport, VPEs do not promote cell survival but are instead involved in cell death through the destruction of the vacuolar membrane and the release of hydrolytic enzymes in the cytoplasm. Further, whether VPEs are specifically targeted and transported by autophagy or randomly sequestered inside autophagosomes during non-selective autophagy remains unknown. In our study, we found that LCAT4 directly interacts with ATG8 (**Fig. 1**), likely allowing its recruitment to the phagophore. Under autophagy induction, LCAT4 is subsequently encapsulated inside the autophagosome and traffics to the vacuolar lumen through the autophagic pathway where it participates in the degradation of autophagic bodies (**Fig. 2, 5**). AlphaFold3 multimer analyses predict that a single AIM domain in LCAT4, located at positions 448 to 451 (YVIL), binds the ADS domain of ATG8 and, accordingly, the ATG8^ADS^ mutant is unable to pulldown LCAT4 *in vitro* (**Fig. 1**). Future analyses of point mutants in the AIM domain of LCAT4 should address the determinants of LCAT4-ATG8 interaction and its relevance for the localization and trafficking of LCAT4. In addition, further work is required to determine the prevalence of autophagy as a transport system for vacuole-bound hydrolases in plants.

In sum, our study unravels the trafficking function of autophagosomes in plants and characterizes a multi-component pathway required for the efficient and specific disruption of membranes of the autophagic bodies, thus highlighting this under-investigated step as a pivot for the completion of autophagy degradation.

## Material and methods

### Arabidopsis lines

All experiments were performed in *Arabidopsis thaliana* of the ecotype Col-0 as wild-type. The following transgenic lines were used as previously described: 35S::GFP-ATG8A (Thompson et al., 2005), pUBQ10::GFP-ATG8A (Shin et al., 2014), pUBQ10::mCherry-ATG8F (Zhuang et al., 2013), YFP-ATG18A (Zhuang et al., 2017), *atg5-1* (Le Bars et al., 2014), *atg7-2* transformed with pUBQ10::GFP-ATG8A (Shin et al., 2014). TDNA insertion lines in *LCAT3 (lcat3;* SALK_035317) and *LCAT4 (lcat4*; SALK_147672.52.80) were obtained from the Nottingham *Arabidopsis* Stock Centre; homozygous lines were obtained by segregation and PCR genotyping using primers listed in Table S1.

LCAT4-tagRFP, LCAT4-tagBFP and LCAT3-GFP plants were generated as following. The *LCAT4* or *LCAT3* coding sequence was PCR-amplified using primers listed in **Table S1** and cloned into the pDONR221 plasmid by Gateway BP reaction. The gene sequences were verified by sequencing. The resulting plasmids were used for multisite gateway LR cloning together with the promoter UBQ10 cloned in the pDONRP4-P1R and tagRFP, tagBFP or GFP cloned in the pDONR2RP3 resulting in the construct pUBQ10-LCAT4-RFP, pUBQ10-LCAT4-BFP and pUBQ10-LCAT3-GFP in the expression plasmid PH7m34GW (LCAT4) or pLOCK180-pFR7m34G (LCAT3). The constructs were verified by sequencing and then transformed into GFP-ATG8A expressing plants for LCAT4-RFP, into YFP-ATG18A expressing plants for LCAT4-BFP and in mCherry-ATG8F for LCAT3-GFP by floral dip. Homozygous lines containing single construct insertions lines were selected based on Hygromycin B resistance (PH7m34GW-LCAT4) or based on red seed coat under epifluorescent microscopy with a set of filters: excitation BP 540-580nm, emission LP 593nm (pLOCK180-pFR7m34G-LCAT3).

To generate *Arabidopsis* lines double knock-out for *LCAT4* and *LCAT3*, *lcat3* and *lcat4* were crossed and the double homozygous line was selected by PCR using primers in **Table S1**. To generate *Arabidopsis* lines expressing pUBQ10::GFP-ATG8A in the different genetic backgrounds, *lcat3*, *lcat4* or *lcat3 lcat4* were crossed with pUBQ10::GFP-ATG8A. The double and triple homozygous lines were selected by Hygromycin B resistance and PCR genotyping using primers listed in **Table S1**.

For the generation of *amiRNA:lcat4* and *amiRNA:lcat3* lines, artificial microRNA sequences specific to the *LCAT4* gene (AT4G19860) or *LCAT3* gene (AT3G03310) were designed using the software WMD3 Web MicroRNA Designer. *amiRNA:lcat4* or *amiRNA:lcat3* precursor fragments were PCR amplified from the pRS300 plasmid as previously described (Schwab et al., 2006) using specific primers listed in **Table S1**. PCR products were cloned into the pENTR/D-TOPO vector (pENTR Directional TOPO Cloning kit, Invitrogen). Clones were verified by sequencing and then transferred by gateway LR cloning into the destination vector PMDC7 which allows expression of the transgene under the control of β-estradiol (Curtis and Grossniklaus, 2003). The constructs were transformed into *Arabidospis* 35S::GFP-ATG8A by floral dip; homozygous lines containing single construct insertions lines were selected based on Hygromycin B and Basta resistance.

### Growth conditions, autophagy induction and chemical treatments

Seeds were vernalized in water and darkness, at 4°C, for 24h to 48h. The seeds were surface sterilized in 10% bleach for 30 min, and sown on Murashige and Skoog (MS) agar medium plates (4.4 g.L^−1^ MS powder including vitamins (Duchefa Biochemie M0222), 0.8 % plant agar (Duchefa Biochemie, P1001), 1% sucrose (Merck Millipore, 84100) and 2.5 mM 2-(N-morpholino)-ethanesulphonic acid (MES, Euromedex EU0033), pH 5.7). Seedlings were grown vertically for 7 days, at 21°C, under long-day conditions (16 h-light/8 h-dark photoperiod, 300 μE m^-^²s^−1^).

To induce autophagy, seedlings were transferred from MS plates to liquid media lacking nutrients as described hereafter for different times indicated in the figures. **(-N):** seedlings were transferred from MS plates into liquid MS-N medium (Murashige and Skoog liquid medium depleted for nitrogen: Murashige and Skoog micronutrient salts (Sigma, M0529), 3 mM CaCl_2_ (Sigma, C5670), 1.5 mM MgSO_4_ (Euromedex, P027), 5 mM KCl (Sigma, P9333), 1.25 mM KH_2_PO_4_ (Sigma P5655), 0.5 % (w/ v) D-mannitol (Sigma M9647), 3 mM MES, pH 5.7). **(-NC):** seedlings were transferred into liquid MS-N medium and incubated in darkness (wrapped in aluminum foil) for different times, as indicated in the figures. As control, for rich conditions (+NC), seedlings were systematically transferred from MS plates to full liquid MS medium (4.4 g.L^−1^ MS powder including vitamins, 1 % sucrose, and 2.5 mM MES, pH 5.7) under normal light conditions. Alternatively, autophagy was induced using the TOR inhibitor AZD-8055 (AZD, MedChemExpress, HY-10422) at concentration ranging from 1 μM (in **Fig. S4**) to 5 μM (in **Fig. S10**) in liquid +N medium for the indicated times. BCECF (B1170, Thermofisher) was used to stain the vacuolar lumen (Krebs et al., 2010). Seedlings were transferred in +N, -NC or +AZD liquid medium for 2 hours prior to the addition of BCECF at 10 µM in the same medium in the dark for 1 hour. To block autophagosome formation, seedlings were treated with Wortmannin (Sigma-Aldrich, 681675; Shin et al., 2014) for 3 hours by transferring plants in liquid -NC medium + 1 μM Wortmannin. Concanamycin A (1 μM; Sigma-Aldrich, C9705) was used as previously described as an inhibitor of vacuole acidification to accumulate autophagic bodies in the vacuole (Huss et al., 2002) by transferring plants in liquid -NC medium + CA for the indicated times. For all chemical treatments, DMSO (dimethyl sulfoxide, Sigma D8418) was used as vector control (untreated plants) in the same conditions (volume, time, type of medium) than the treated plants. To induce the expression of the artificial micro-RNA against *LCAT4* or *LCAT3*, seedlings were transferred in liquid medium containing 10 μM of β-estradiol (Sigma Aldrich, E2758) prior to autophagy induction. The incubation time varied from 8 to 16 hours according to the experiments and are indicated in the respective figures.

### Structural analysis using AF3

AlphaFold 3 (https://www.alphafoldserver.com), developed by DeepMind, is an advanced AI platform for predicting protein structures. Protein sequences analyzed in this study were obtained from UniProt. The selected sequences were then submitted to the AlphaFold prediction module for detailed structural evaluation. The align command in UCSFChimera (version 1.17) was employed to superimpose the AF3 predictions onto known structures and to show the confidence score of the AF3 predictions using the local distance difference test (pLDDT) scores on the lDDT-Ca metric. pLDDT corresponds to the model’s predicted score on the lDDT-Cα metric (Ibrahim et al., 2023) and PAE plots provide valuable information about the reliability of relative position and orientations of different domains. PAE heatmap were designed using https://subtiwiki.uni-goettingen.de/v4/paeViewerDemo (Elfmann and Stülke, 2023).

#### *In vitro* immunoprecipitation

All recombinant proteins were produced using *E. coli* strain Rosetta2 (DE3) pLysS. AIM wt or AIM mut were used as previously described (Stephani et al., 2020). Transformed cells were grown in 2xTY media supplemented with 100 μg/mL Spectinomycin at 37°C to log phase (OD_600_ 0.6–0.8), followed by induction with 300 μM isopropyl β-D-1-thiogalactopyranoside (IPTG) and incubation at 18°C overnight. Cells were harvested by centrifugation for 10 minutes, 4,000 rpm at 4°C, supernatant was discarded, and pellets were stored at -20°C until further use. Bacterial pellets were resuspended in 5 mL Resuspension Buffer (100 mM NaPi pH 7.2, 300 mM NaCl, 1 mM DTT) supplemented with 1x cOmplete Protease Inhibitor Cocktail tablet (Roche) and 1 μg/mL Benzonase. Cells were lysed twice by 1-3 seconds on-off sonication cycles at 30% amplitude (Vibra Cell VCX750, Sonics & Materials) and cleared by centrifugation for 30 minutes, 20,000 rpm at 4°C in a top-table centrifuge. 5 µl of glutathione magnetic agarose beads (Pierce Glutathione Magnetic Agarose Beads, ThermoFisher Scientific) were equilibrated by washing them with IP buffer (100 mM NaPi pH 7.2, 300 mM NaCl, 1 mM DTT, 0.01% (v/v) IGEPAL® CA-630). Normalized *E. coli* clarified lysates were mixed and the final reaction volume was brought to 1 mL with IP Buffer. Reactions were added to the washed beads and incubated on an end-over-end rotator for 1 hour at 4°C. After incubation, beads were washed 5 times in 1 mL wash buffer. Bound proteins were eluted and denatured in 50 µL 2X SDS Laemmli buffer (2% SDS, 10% glycerol, 5% 2-β-mercaptoethanol, 0.002% Bromophenol blue and 0.0675 M Tris HCl, pH 6.8) for 5 minutes at 95°C prior to Western blotting. For Western blotting, the indicated total protein amount was loaded on 4-20% Mini-PROTEAN TGX precast SDS-PAGE gels (Bio-Rad) and blotted on nitrocellulose membranes (Bio-Rad) using the semi-dry Trans-Blot Turbo Transfer System (Bio-Rad). Membranes were blocked in TBS-T (10 mM Tris-HCl pH 7.5, 150 mM NaCl, 0.1% Tween 20) + 5% skimmed milk at room temperature for 1 hour. After blocking, membranes were incubated with the respective primary antibody diluted in blocking buffer either for 1 hour or overnight at 4°C. The primary antibody, an anti-MBP (Sigma Aldrich, # M1321) at a dilution of 1/5000, was recovered, and membranes were washed 5 times for 5 minutes with TBS-T before incubation with the respective secondary antibody, an Anti-Mouse IgG-HRP Conjugate (Bio-Rad, #1706516) at 1/5000 or Anti-GST HRP-Conjugate (GE Healthcare, # RPN1236) at 1/5000, for 45 minutes at room temperature. The immune reaction was developed using Pierce™ ECL Western Blotting Substrate (ThermoFisher) and detected with iBright Imaging System (Invitrogen).

### Phenotypic assays

For germination assays, seeds were vernalized for two days in water prior to being sown on MS ½ + 1% sucrose. 20-30 seeds of each genotype were analyzed per plate for a total of 60 seeds per genotype. Plates were placed horizontally at 21°C, under long-day conditions (16 h-light/8 h-dark photoperiod, 300 μE m^-^²s^−1^). At the indicated times, seed germination was assessed qualitatively by examining the tegument of the seed and the emergence of the radicle. Results are expressed as the percentage of germinated seeds over total seeds per independent plate. The experiment was performed in two biological replicates with lot#1 (each genotype in the Col-0 background) and lot#2 (each genotype expressing pUBQ10:GFP-ATG8A). For root length measurement, similar settings and conditions were used for plant growth *in vitro*. Each plate was imaged at 54 h, 72 h and 96 h after sowing and root length was marked. Root length was measured using the Fiji software (National Institutes of Health, USA, http://imagej.nih.gov/ij) using the aforementioned images. For comparison of vegetative growth and onset of senescence, seedlings of each indicated genotype were sown on 0.8 % plant agar plates containing full-MS powder including vitamins, 1% sucrose and 2.5 mM MES and then grown at 21°C under an LD (16-h-light/8-h-dark) photoperiod. After 1 week, seedlings were transplanted to soil and grown at 21°C under SD (8-h-light/16-h-dark photoperiod) conditions for either 6 or 12 weeks and imaged. To assess the effects of nutrient limitation on plant physiology, seedlings of each indicated genotype were grown for 1 week on MS plates in rich conditions as described above. Seedling were then transferred for 10 days to a solid minimal medium lacking sucrose and nitrogen (medium was prepared as in Gaude et al., 2007, without sucrose) and wrapped with aluminum foil. After 10 days, plates were imaged and the color of the seedlings was assessed qualitatively. Nine independent plates containing about 20 seedlings of each genotype, grown and treated side by side, were analyzed. For each plate, the percentage of green seedlings was calculated as the number of mostly green seedlings over total seedlings.

### Microscopy analysis

All live confocal observations were performed on root epidermal cells of the transition zone and elongation to early differentiation zone, employing 7-day-old seedlings mounted in culture medium as indicated in the figures. Roots were kept under the microscope for no longer than 10 min in order to avoid secondary effects due to prolonged treatment times. Confocal images were acquired with a ZEISS LSM880 confocal system. Laser excitation lines for the different fluorophores were 405 nm for BFP, 488 nm for GFP, 514 nm for YFP and 561 nm for RFP or mCherry. Fluorescence emissions were detected at 420-470 nm for BFP, 490–597 nm for GFP, 520-555 nm for YFP, 580–650 nm for RFP and 561-633 nm for mCherry. In multi-labeling acquisitions, detection was in sequential line-scanning mode. Confocal microscope images were processed using the Zen Black Software (Zeiss) for intensity optimization. For puncta quantification and co-localization analyses in the cytosol, n number of images as indicated in the figure legends were taken in a minimum of 5 independent roots in B number of biological experiments as indicated in the figure legends. Puncta were counted manually in each image and each channel and root surface areas were quantified using Fiji to obtain the puncta/surface values. When multiple z-planes were imaged, the number of puncta and the surface of the area was quantified in each plane and averaged. For puncta quantification and co-localization analyses in the vacuole, a similar image acquisition and analytical pipeline protocol was followed with the exception that puncta were counted manually in selected cells where the vacuole was clearly discernable from cortical cytosol (using the GFP-ATG8A signal as a marker of the cytosol). The surface of each analyzed cell was quantified to obtain the puncta/surface values. Co-localization between ATG8 and LCAT3 or LCAT4 was assessed manually comparing the position of puncta in each channel, overlapping signals indicated colocalization. Co-localization was confirmed on selected puncta, by comparing the maximum intensity of signal in each channel plotted along a line using the plot profile tool of Fiji. To measure the vacuole/cytosol signal intensity ratio in **Fig. 2c**, cells (number indicated in the figure legend) where the vacuole was clearly discernable were selected in 5-11 independent roots. Signal intensity was quantified using Fiji, either at the cell periphery or inside the vacuolar lumen, in a delimitated region of interest of the same surface for each location. In **Fig. 4c**, high-zoom time series images of vacuoles were analyzed using the TrackMate plugin (Ershov et al., 2022). Tracking was done on the mCherry-ATG8 channel, while LCAT3 co-localization events were manually identified. Images of localization of CPY-LCAT3-GFP and CPY-LCAT4-GFP in yeast were acquired using a CCD cool Snap HQ2 camera mounted on a ZEISS AxioImager epifluorescence microscope. GFP was observed using a dedicated filter setup (Excitation 472/30, Emission 520/55).

### Western-blot analyses

To assess autophagy flux, we performed the GFP-ATG8 processing assay: 7-day-old seedlings were transferred from full MS plates into rich MS liquid medium (+NC), -N liquid medium (-N) or −NC liquid medium (-NC) and total proteins were extracted after treatments. Whole seedlings were frozen in liquid nitrogen, disrupted using a TissueLyser (Qiagen), and homogenized in the following buffer: 100 mM Tris pH 7.5, 200 mM NaCl, 1 mM EDTA, 2% β-mercaptoethanol, 0.2 % Triton 100X, 1 mM PMSF and antiprotease mix (P9599 Sigma). Homogenates were centrifuged at 1600 × g for 20 min at 4°C and twice at 1600×g for 10 min at 4°C, transferring only the supernatant at each stage. The protein concentration of each sample was determined using Bio-Rad Protein Assay Dye Reagent Concentrate, (BioRad, ref #5000006) and measuring sample absorption at 595 nm. Equal amounts of proteins were denatured using Laemmli buffer at 55 °C for 15 min and loaded on acrylamide gels. After migration on 12% SDS-PAGE (TGX Stain-Free FastCast Acrylamide kit, BioRad), equal protein quantity per lane was verified by stain-free activation of the loaded gel. After transfer to nitrocellulose (BioRad, #1704270), membranes were incubated with an anti-GFP mouse antibody (Roche, #11814460001) at a dilution of 1/10,000 for 35S:GFP-ATG8A expressing plants (**Fig. S6, S9**). For pUBQ10:GFP-ATG8A plants, blots were cut to separate the GFP-ATG8 bands (dilution at 1/5,000) from the GFP bands (dilution at 1/10,000) to allow quantitative analyses of each band without saturation of the free GFP signal (**Fig. 3-5**). Peroxidase activity coupled to the Goat anti-Mouse antibody at a dilution of 1/10,000 (Bio-Rad, #1706516) was revealed using Clarity Max Wester ECL (Biorad #170562) and detected with the Chemidoc MP Imaging system (Bio-Rad). Blots were imaged at different exposition times, long enough to observe clear distinct bands but prior to pixel saturation. Blot images were analyzed using the Fiji software; band intensities were calculated using the plot surface areas functionality of the software. The intensities of the free-GFP band and the GFP-ATG8 band were then used to calculate the relative GFP/GFP-ATG8 ratios and compare it to that of WT which was set to 1 in each independent experiment.

To assess the level of LCAT4 upon nutrient starvation, 7 day-old plants expressing pUBQ10::LCAT4-RFP were transferred in (+N) or (-NC) liquid medium for 3 hours. Total proteins were extracted, quantified and denaturated as describe above. Equal amounts of total proteins were loaded on 12 % SDS-PAGE gels and the level of LCAT4-RFP was analyzed by immunoblotting using anti-RFP antibody (Abcam, ab34767; 1/1000). Peroxidase activity coupled to the Goat anti-mouse antibody at a dilution of 1/10,000 (Bio-Rad #1706516) was used for revelation. Equal protein quantity per lane was verified by stain-free activation of the loaded gel. Blot images were analyzed using the Fiji software as described above and the amount of LCAT4 was normalized to that of total protein measured in the stain free.

### RT-QPCR

Total RNA was extracted using the RNeasy mini kit (Qiagen). To eliminate genomic DNA contamination, an additional DNase treatment was performed according to the RNeasy kit instruction with the DNA removal kit (Invitrogen). One microgram of total RNA was reverse transcribed into cDNA in a 20μL reaction mixture using the Superscript IV reverse transcriptase enzyme (Invitrogen). cDNA levels were then analyzed using the iQ™ Sybr Green supermix (BioRad) on the iQ iCycler thermocycler (BioRad) with the gene-specific primers listed in **Table S1**. The thermocycling program consisted of one hold at 95°C for 3 min, followed by 40 cycles of 15 s at 95°C and 30 s at 58°C, and a melt curve from 65°C to 90°C with an increment of 0,5°C all 5s. After completion of these cycles, melting-curve data were then collected to verify PCR specificity, contamination, and the absence of primer dimers. The transcript abundance in samples was determined using a comparative threshold cycle method. The relative abundance of the reference mRNAs of *ACTIN 2/8*, *AT4G33380* or *SAND* was determined in each sample and used for normalization, according to the experiment.

### Yeast strains, expression, media and culture

Yeast strains used in this study are listed in **Table S2**. To generate the PYES3-LCAT3, PYES3-LCAT4 and pYES2-LCAT4 plasmids, the coding sequences of the corresponding genes were cloned from the pDONR221 (see *Arabidopsis* lines) into the PYES3 or pYES2 plasmid (both harboring the GAL1 promoter and either TRP or URA auxotrophic genes, respectively) using LR gateway cloning. To generate pYES3-CPY-LCAT3, pYES3-CPY-LCAT4 or pYES2-CPY-LCAT4, a DNA fragment encoding the first 50 amino acids of the CPY protein (CPY^1-50^; from the *PRC1* gene) was PCR amplified from *S. cerevisiae* total genomic DNA using the primers listed in Table S1. The forward primer was flanked with the Attb1 sequence while the reverse primer was flanked with a DNA fragment consisting of the first 26 nucleotides of *LCAT4* or first 20 nucleotides of *LCAT3*. The coding sequence of *LCAT3* and *LCAT4* were PCR amplified using primers listed in **Table S1**, with the forward primer flanked with the last 24 nucleotides of CPY^1-50^ and the reverse primers flanked with Attb2 sequence. Then, the CPY^1-50^ fragment and LCAT3 or LCAT4 coding sequence were used as template and fused using overlapping PCR to generate CPY^1-50^-LCAT3 or CPY^1-50^-LCAT4. The resulting DNA fragments were cloned into the PDONR221, sequence verified and then transferred into the PYES3 or pYES2 plasmid using gateway cloning. pYES3-CPY-LCAT3^S177A^ was generated similarly after performing site directed mutagenesis to change serine 177 to an alanine by overlapping PCR using primers listed in Table S1.

**In Fig.6a-c**, to express LCAT3, LCAT4, CPY-LCAT3, CPY-LCAT4 or CPY-LCAT3^1-50^, SSY12 yeast cells (see **Table S2**) were transformed with the corresponding plasmids and transformants were selected on synthetic minimal medium lacking tryptophane (SMD; 0.67 % yeast nitrogen base, 2 % glucose, 0.2 % drop out -Trp). As controls, SSY12 or SEY6210 yeast cells were transformed with the empty PYES3 plasmid. To induce the expression of the transgenes, yeast cells were grown overnight in gal-SMD lacking tryptophane (0.67 % yeast nitrogen base, 2 % galactose, 0.2 % drop out). The following morning, cultures were diluted back to 0.2 OD and yeast cells were grown until mid-log phase (OD 0.6-1). 1 OD of each culture was collected and total protein were extracted as previously described (*Bernard et al., 2015)*. Equal amounts of proteins were loaded on 12% SDS-PAGE gels and immunoblot analyses were performed using nitrocellulose membranes, the Ape1 primary antibody (1/10,000; Klionsky et al., 1992) and a peroxidase-coupled Goat anti-Rabbit secondary antibody (1/10,000; Biorad, #1706515).

In **Fig. 6d-g**, to construct GFP-AtATG8A expressing cells, the coding sequence of GFP and that of AtATG8A were PCR amplified and fused using overlapping PCR using primer listed in **Table S1**. The resulting transgene was digested with PacI and AscI and ligated into the pFa6a-HIS plasmid (Longtine et al., 1998). Endogenous Atg8 was disrupted and replaced by GFP-AtATG8A-HIS at the Atg8 locus using a standard method for chromosomal tagging (Longtine et al., 1998) by amplifying GFP-AtATG8A-HIS from the previously constructed pFa6a:GFP-AtATG8A-HIS plasmid using primers listed in **Table S1** and transforming either WT cells (SEY6210) or *atg15Δ* cells (TKYM108), see Table S2. Transformants were selected on minimal medium lacking histidine (SMD; 0.67 % yeast nitrogen base, 2 % glucose, 0.2 % drop out -His). Replacement of endogenous Atg8 by GFP-AtATG8A was verified by PCR using primers listed in **Table S1**. The resulting WT cells (YAB610) or *atg15Δ* cells (YAB613) expressing GFP-AtATG8A were used for further yeast transformation to express either CPY-LCAT3 (cloned in pYES3), LCAT4 or CPY-LCAT4 (cloned in pYES2) and a combination of CPY-LCAT3 (cloned in pYES3) and LCAT4 (cloned in pYES2) or empty plasmids as listed in **Table S2**. Yeast cells transformed with pYES3 plasmids were selected on minimal medium lacking tryptophane; cells transformed with pYES2 plasmids were selected on minimal medium lacking uracil and cells transformed with pYES2 and pYES3 plasmids were selected on minimal medium lacking both tryptophane and uracil (SMD; 0.67 % yeast nitrogen base, 2 % glucose, 0.2 % appropriate drop out). To induce the expression of the transgenes, yeast cells were grown overnight in gal-SMD lacking either tryptophane, uracil or both (0.67 % yeast nitrogen base, 2 % galactose, 0.2 % drop out). The following morning, cultures were diluted back and yeast cells were grown until mid-log phase (OD 0.6-1) prior to inducing autophagy by transferring cells into SD-N medium (0.17% yeast nitrogen base without amino acids, containing 2% galactose). In **Fig. 6d,f**; cells were starved for 4 hours; in **Fig. 6e,g**, cells were starved for 10 hours. 1 OD of each culture in +N or -N condition was collected. Protein extraction, immunoblot and GFP-ATG8 processing were performed as previously described (Bernard et al., 2015). Equal amounts of proteins were loaded on 12% SDS-PAGE gels and immunoblot analyses were performed using nitrocellulose membranes, the GFP primary antibody (1/10,000; Roche, #11814460001) and a peroxidase-coupled Goat anti-Mouse secondary antibody (1/10,000; Bio-Rad #1706516).

### Statistical analyses

Statistical analyses were performed using GraphPad Prism 9.3.1 (GraphPad Software, La Jolla, CA, USA) with tests indicated in the figure legends. P-values are as follows: P-value > 0.05 (non-significant, ns), *P <0.05, **P <0.01 and ***P < 0.001.

## Acknowlegments

We thank Professor L. Jiang (The Chinese University of Hong Kong) for YFP-ATG18A seeds and Professor Taijoon Chung (Pusan National University, Korea) for pUBQ10:GFP-ATG8A seeds.

## Author contributions

JC designed and performed experiments and wrote the manuscript. MB, VSMH, TI, EAM designed and performed experiments and participated in the redaction of the manuscript. ITM, REG, FDD, JL, CC, SP, FD, JJ performed experiments and participated in the redaction of the manuscript. TOB, YD designed experiments and participated in the redaction of the manuscript. AB designed research, performed experiments, wrote and edited the manuscript.

## Funding

This project has received funding from the European Research Council (ERC) under the European Union’s Horizon 2020 research and innovation program (grant agreement No 852136 to AB) and from Idex Bordeaux (AutoLip Emergence Program to AB). Microscopy was done at the Bordeaux Imaging Center, a member of the national infrastructure France-BioImaging supported by the French National Research Agency (ANR-10-INBS-04).

### Declaration of interests

The authors declare no competing interests.

**Supplementary Figure 1. LCAT4 co-localizes with YFP-ATG18A.**

Representative confocal images of 7-day-old seedlings co-expressing YFP-ATG18A and LCAT4-BFP. Plants were placed in liquid MS medium in rich condition (+NC) or deprived of nutrients for 1h (−NC 1h). Arrowheads indicate colocalization. Scale bar, 10 μm.

**Supplementary Figure 2. Prediction of the LCAT4-ATG8 interaction**

**(a)** Analysis of the amino acid sequence of LCAT4 shows several [W,Y,F][X][X][L,I,V] motifs, highlighted in grey. The AlphaFold3 predicted AIM domain ^448^YVIL^451^ is highlighted in orange. (**b-e**) Predicted aligned error (PAE) plots and graphic representation of three models based on AlphaFold3 multimer modeling of the LCAT4-ATG8 interaction using WT LCAT4 (b,c) or a mutated version of LCAT4 where the four residues of the predicted AIM domain where replaced by alanines (^448^YVIL^451^_AAAA, d,e). Units: amino acid residues; light green to dark green: expected position error in angstroms. IpTM are indicated at the top of each PAE plots. Three best-scoring models from AlphaFold3 show almost identical binding interfaces between WT LCAT4 and ATG8 (b,c). The LCAT4-ATG8 interaction interface is shown by the arrows in each protein (b). Neither of the three models from AlphaFold3 show an interface between LCAT4^448^YVIL^451^_AAAA and ATG8 (d,e).

**Supplementary Figure 3. LCAT4 binds all ATG8 isoforms *in vitro* in an AIM-dependent manner.**

**(a)** LCAT4 binds all ATG8 isoforms *in vitro*. Bacterial lysates containing recombinant protein were mixed and pulled down with glutathione magnetic agarose beads. Input and bound proteins were visualized by immunoblotting with anti-GST and anti-MBP antibodies. **(b)** LCAT4 interacts with ATG8 in an AIM-dependent manner. ATG8A^ADS^=ATG8A^(Y50A,L51A)^. AIM *wt* and AIM *mut* peptides were added to a final concentration of 200 µM. Bacterial lysates containing recombinant protein were mixed and pulled down with glutathione magnetic agarose beads. Input and bound proteins were visualized by immunoblotting with anti-GST and anti-MBP antibodies.

**Supplementary Figure 4. LCAT4 is found in the vacuole under autophagy induction conditions.** Representative confocal images of Arabidopsis roots expressing LCAT4-RFP in the transition zone. Plants were imaged in nutrient rich condition (+NC), after 3 hours in nutrient starvation (-NC) or +NC with the addition of AZD8055 (AZD; 1 μM). Vacuoles were stained with BCECF/AM (BCECF, 10 µM). Scale bar, 10 µm.

**Supplementary Figure 5. Characterization of the *LCAT4* and *LCAT3* transgenic lines**

**(a)** Representative schemes of t-DNA insertion in the *LCAT4* or *LCAT3* genes. Exons, thick squares; introns, thin lines. Positions of PCR primers for genotyping are indicated; gene-specific (LP, RP) and tDNA specific (RB). **(b)** PCR genotyping of *lcat3*, *lcat4* and *lcat3,4* mutants compared to WT and *atg5* plants*. ECR* was amplified as control. **(c)** *LCAT4* and *LCAT3* mRNA levels in *lcat3*, *lcat4* and *lcat3,4* mutants compared to WT (set to 1 in each experiment) in 7 -day-old seedlings. Results represent the average ± SD of 3 replicates; normalized by the reference genes *SAND* and *ACTIN 2/8*. **(d,e)** Analyses of the transcripts in *amiRNA:LCAT4* (d) or *amiRNA:LCAT3* (e) lines showing a decrease in the expression of the target gene compared to WT plants. 7-day-old seedlings were treated with β-estradiol (10 µM) for 24 h in (d) or 8 h in (e). Results represent the average ± SD of 3 replicates; normalized by the reference genes *ACTIN 2/8* and *AT4G33380* in (d) or by *ACTIN 2/8* in (e).

**Supplementary Figure 6. Analysis of the autophagic flux in the *amiRNA:LCAT4 line*.**

**(a)** Immunoblot analyses of GFP-ATG8A and its degradation product in WT or *amiRNA:LCAT4* plants expressing 35S:GFP-ATG8A. Seedling were transferred to either (+N) or (-NC) for 16 hours +/-10 µM of β-estradiol. Total proteins were extracted and analyzed with anti-GFP antibodies. Stain free images were used as loading control. **(b)** Quantification of the ratio of GFP/GFP-ATG8A in (a) relative to that of WT plants in (-NC) condition which was set to 1 in each experiment. Results are presented as the average with values of distinct replicates; n=4 biological replicates.

**Supplementary Figure 7. LCAT1 and LCAT3 are homologs of LCAT4**

**(a)** Phylogenetic tree of *Arabidopsis* phospholipases. The amino acid sequences were aligned using the Clustal Omega algorithm; phylogenetic analyses were performed using MEGA11. Phylogenetic relationships between taxons were established using the maximum likelihood method and the robustness of the tree obtained was tested with a bootstrap of 1000 replicas. Percentage of identity between LCAT4 and the other LCAT members is indicated in green. **(b)** Relative abundance of the transcripts of *LCAT1, LCAT3* and *LCAT4* in 7-day-old seedlings placed in rich condition (+N, liquid medium) or deprived of nutrients (-NC, liquid medium) for 6h. Levels of mRNA were normalized by the reference genes *ACTIN 2/8*, *AT4G33380* and *SAND* and compared to +NC conditions which were to set to 1 in each experiment. Results present the average of ± SD of 3 replicates each in two independent experiments (-NC#1; - NC#2).

**Supplementary Figure 8. Comparison of the protein and models of LCAT3 and LCAT4 suggest that LCAT3 does not interact with ATG8**

**(a)** Alignment between the amnio acid sequence of LCAT4 (magenta) and LCAT3 (grey). The amino acids of LCAT3 identical to that of LCAT4 are indicated in magenta. The predicted AIM domain of LCAT4 is depicted in orange. The C-terminal tails of the protein are delineated by a yellow dotted square in (a) and (b). **(b)** AlphaFold3 models of LCAT3 and LCAT4 shows major differences in the C-terminal tail of the two proteins (yellow dotted rectangle). **(c,d)** Modelling the interface between LCAT3 and ATG8 did not result in a single orientation of ATG8 towards LCAT3 as shown for three best-scoring models calculated by AlphaFold3 multimer (c). Predicted aligned error (PAE) plots of three models of the LCAT3-ATG8 predict no interaction between the proteins (d).

**Supplementary Figure 9. The inducible knocking down of *LCAT3* causes a slight reduction in autophagy flux in -N conditions but not in -NC.**

**(a)** Representative image of GFP-ATG8A processing assay in *amiRNA:LCAT3* compared to WT plants. 7-day-old seedlings were transferred to nutrient rich liquid medium (+N) supplemented with 10µM of β-estradiol for 16 hours to induce the expression of the amiRNA. Seedlings were then transferred to either nutrient rich liquid medium (+N), liquid medium depleted in nitrogen (-N) or depleted in nitrogen and carbon (-NC, in the dark) during 8h, all media containing 10 µM of β-estradiol. For heat stress (H), plants were transferred to +N liquid medium for 5 h at room temperature followed by 3 h at 37°C. **(b)** Quantification of the ratio of GFP/GFP-ATG8A in (a) in percentage of the WT in -NC condition which was set to 1 in each experiment. Results are presented as the average ± SD with values of all replicates collected in three independent biological experiments; in +N, n=3, in -N and -NC, n=6, in H, n=3. Statistical differences were assessed using Mann Whitney test compared to WT in the same condition (+N, -N, H) and one sample t.test compared to WT in the same condition (-NC); p values are indicated.

**Supplementary Figure 10. Knocking out *LCAT3* and *LCAT4* results in the accumulation of autophagic bodies.**

**(a)** Comparative confocal image analysis of 7-day-old seedling roots between *lcat3,4* line and wild-type (WT). Plants were transferred into 6-well plate with 3 ml of nutrient rich liquid complemented with nothing (Mock), AZD8055 (AZD, 5µM) or AZD + concanamycine A (AZD + ConA; 1 μM). Filled arrow heads indicate the autophagic bodies and hollow arrow heads the autophagosomes. Scale bars, 20 µm. **(b)** Quantification of GFP puncta in (a). Results are presented as the number of puncta per 10 μm² of root area and show the average and individual values, from left to right, n=175 pictures of 13 roots, n=190 pictures of 12 roots, n=220 pictures of 14 roots, n=180 of 11 roots, n=185 of 12 roots and n=182 of 12 roots in 3 independent biological replicates.

**Supplementary Figure 11. Knock out mutants for *LCAT3*, *LCAT4* or both *LCAT3,4* show similar growth and development compared to WT plants.**

**(a)** LCAT3 and LCAT4 are not required for proper seed germination while mutants with a complete block in the autophagy pathway (*atg5*, *atg7*) show germination impairments. 20-30 seeds per plate were sown on ½ MS + sucrose after two days of vernalization. Germination was assessed at the indicated times points. Results are expressed as percentage of germinated seeds relative to total seeds sown in 2-3 independent plates. Results of two biological experiments are presented (lot#1, lot#2). A total of 60 seeds were analyzed by independent experiment. **(b)** LCAT3 and LCAT4 knock out mutants show no to little changes in root elongation compared to WT plants and *atg* mutants. Roots were measured at the indicated times after seed sowing. Results are presented as box plots with average, min and max of 2-3 independent plates with a total of 60 seedlings per genotype per independent experiment. Results of two biological experiments are presented (lot#1, lot#2). **(c,d)** Representative image of the vegetative growth and onset of senescence of *lcat3*, *lcat4* and *lcat3,4* mutants compared to WT plants and *atg* mutants. Plants were grown on soil at 21°C under short-day conditions for 6 weeks (c) or 12 weeks (d). Scale bar: 1 cm.

**Supplementary Figure 12. Localization of CPY-LCAT3-GFP and CPY-LCAT4-GFP expressed in *atg15Δ* yeast cells shows signal in the vacuole.** Representative image of n=4 independent replicates. Scale bar, 10 µm.

**Supplementary Movie 1. LCAT3 can transiently associate with autophagosomes close to the vacuole.** Movie representing LCAT3-GFP puncta associating with an autophagosome labeled with mCherry-ATG8F in the cytosol, at the edge of the vacuole. 7-day-old seedlings expressing both transgenes as in Fig. 4 were imaged by confocal microscopy after 3 hours of nutrient starvation with the addition of concanamycin A (-NC 3h+CA, 1 μM). Scale bar, 2 µm.

