## Supplemental figures for "A dual component system instructs membrane hydrolysis during the final stages of plant autophagy"

### Figure S1

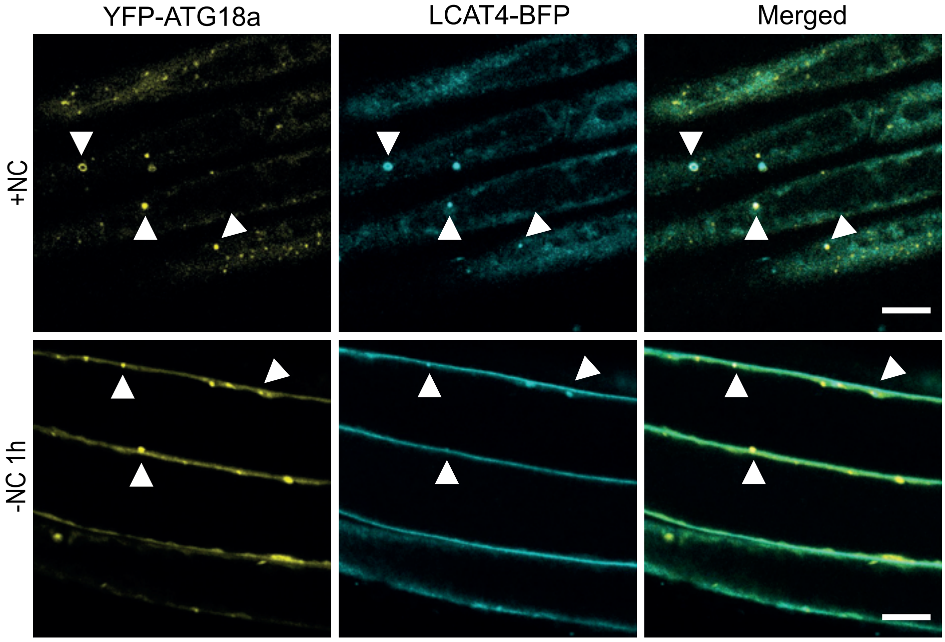

### Figure S2

**a**

```
MSLLLEEIIRSVEALLKLRNRNQEPYVDPNLNPVLLVPGIAGSILNAVDHENGNE
ERVWVRIFGADHEFRTKMWSRFDPSTGKTISLDPKTSIVVPQDRAGLHAIDVLDP
DMIVGRESVYYFHEMIVEMIGWGFEEGKTLFGFGYDFRQSNRLQETLDQFAKKLE
TVYKASGEKKINVISHSMGGLLVKCFMGLHSDIFEKYVQNWIAIAAPFRGAPGYI
TSTLLNGMSFVNGWEQNFFVSKWSMHQLLIECPSIYELMCCPYFKWELPPVLELW
REKESNDGVGTSYVVLESYCSLESLEVFTKSLSNNTADYCGESIDLPFNWKIMEW
AHKTKQVLASAKLPPKVKFYNIYGTNLETPHSVCYGNEKMPVKDLTNLRYFQPTY
ICVDGDGTVPMESAMADGLEAVARVGVPGEHRGILNDHRVFRMLKKWLNVGEPDP
FYNPVNDYVILPTTYEFEKFHENGLEVASVKESWDIISDDNNIGTTGSTVNSISV
SQPGDDQNPQAEARATLTVQPQSDGRQHVELNAVSVSVDA
```

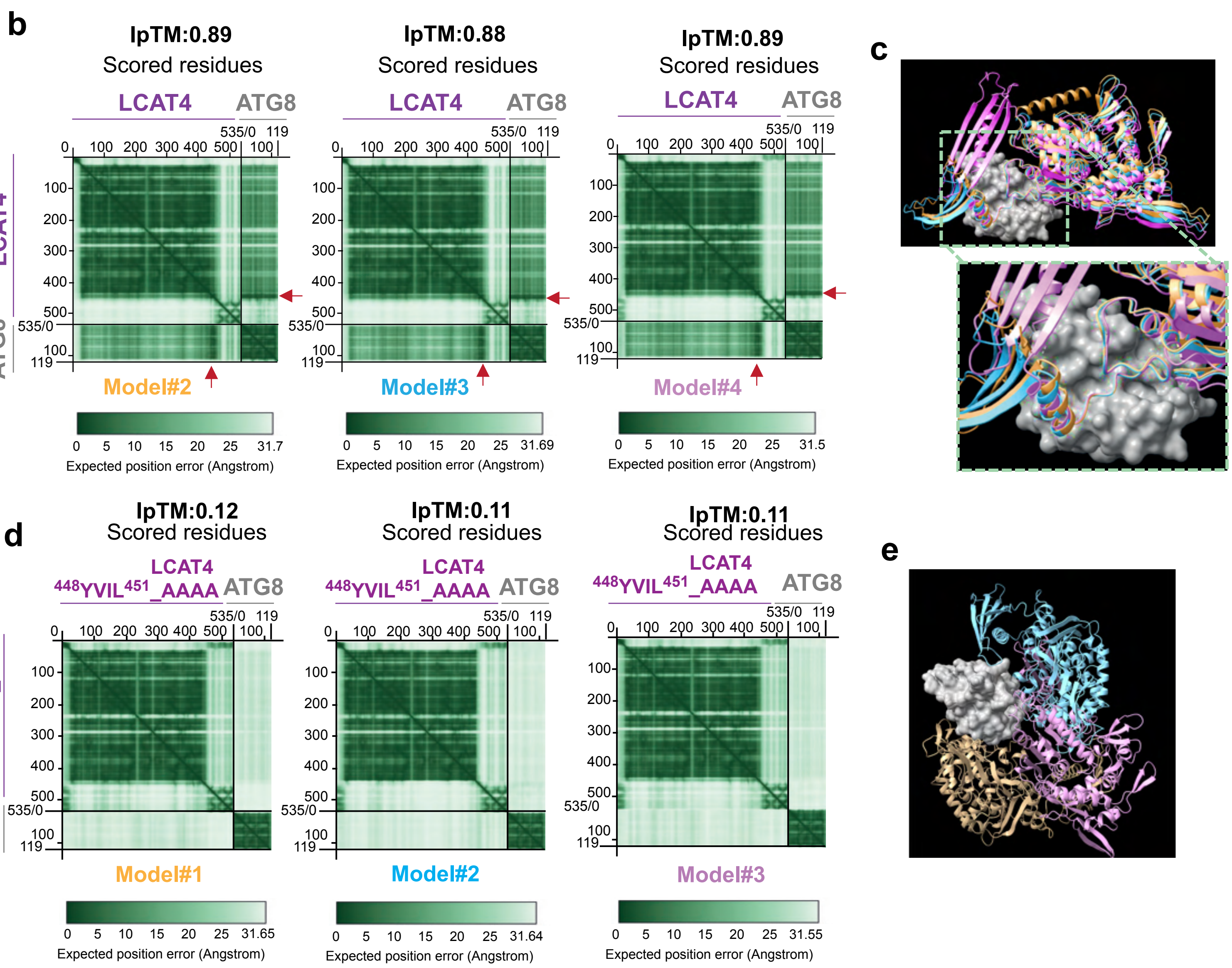

### Figure S3

a

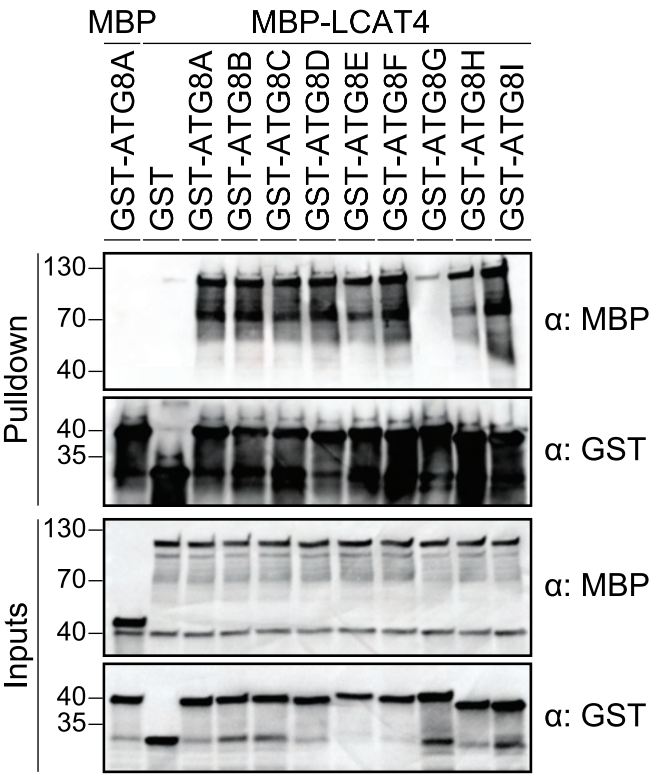
MBP
MBP-LCAT4
GST-ATG8A
GST
GST-ATG8A
GST-ATG8B
GST-ATG8C
GST-ATG8D
GST-ATG8E
GST-ATG8F
GST-ATG8G
GST-ATG8H
GST-ATG8I
Pulldown
Inputs
130
70
40
35
α: MBP
α: GST

b

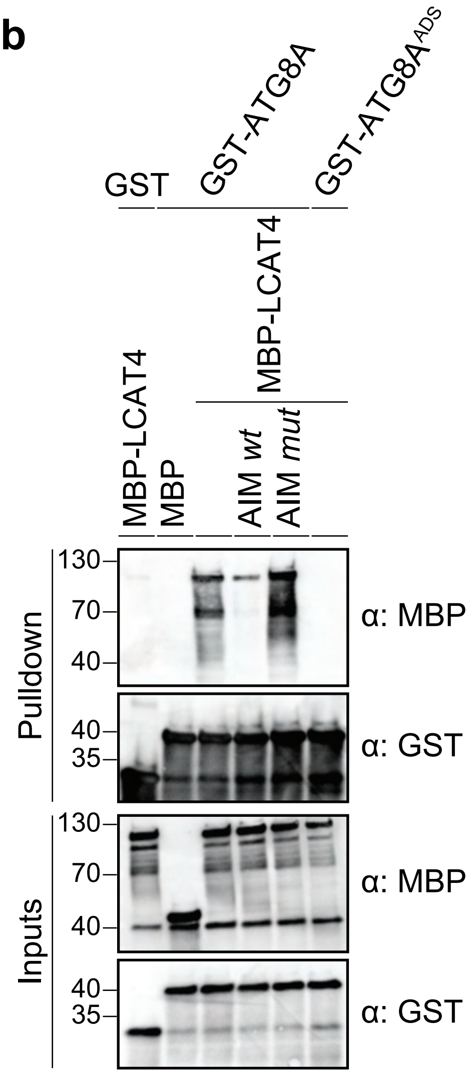
GST
GST-ATG8A
GST-ATG8A^ADS
MBP-LCAT4
MBP-LCAT4
MBP
AIM wt
AIM mut
Pulldown
Inputs
130
70
40
35
α: MBP
α: GST

### Figure S4

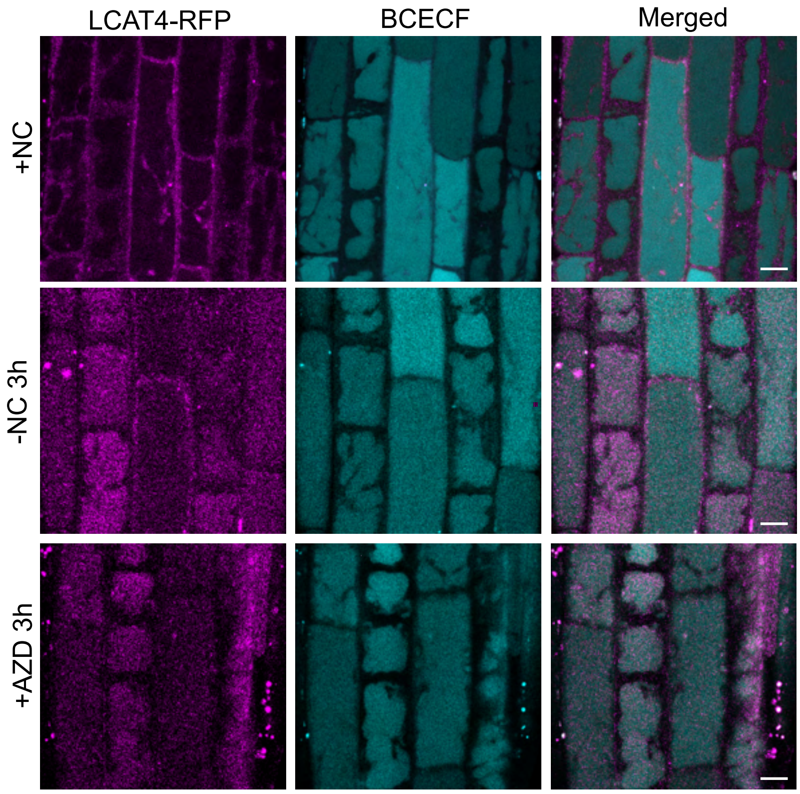

### Figure S5

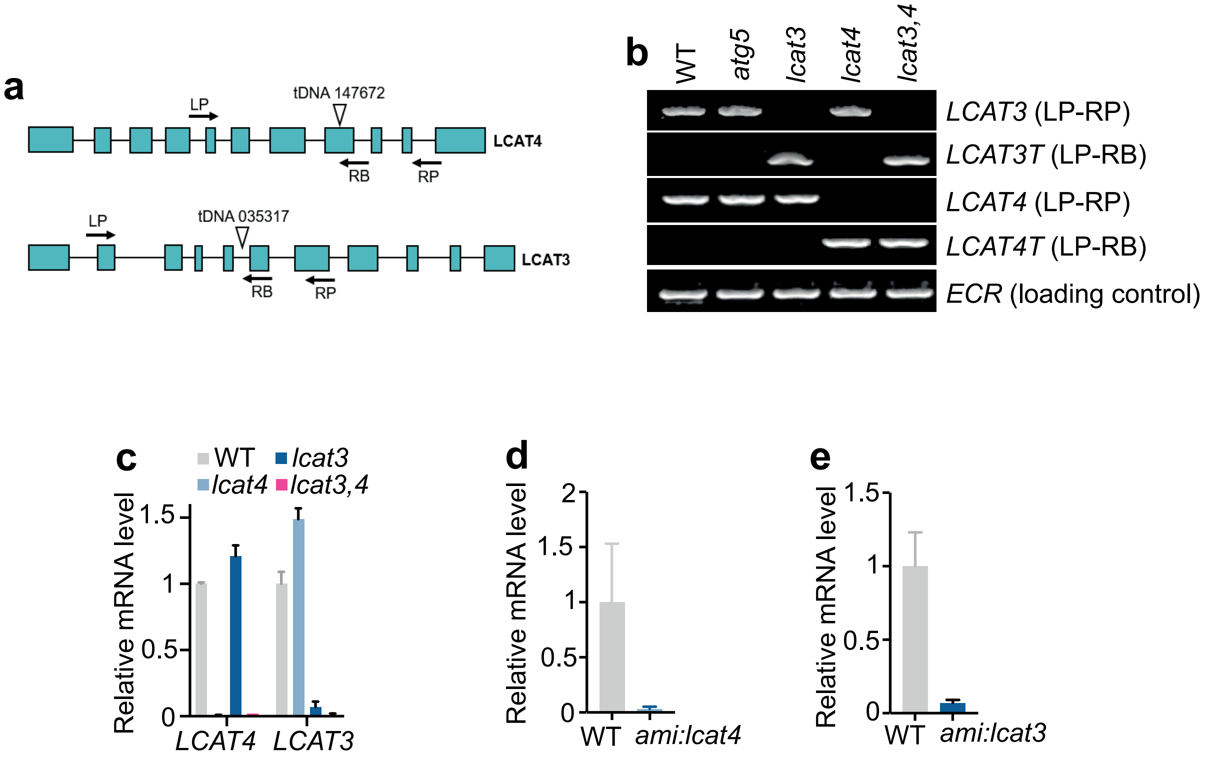

a
LP
tDNA 147672
LCAT4
RB
RP
LP
tDNA 035317
LCAT3
RB
RP
b
WT
atg5
lcat3
lcat4
lcat3,4
LCAT3 (LP-RP)
LCAT3T (LP-RB)
LCAT4 (LP-RP)
LCAT4T (LP-RB)
ECR (loading control)
c
WT
lcat3
lcat4
lcat3,4
Relative mRNA level
1.5
1
0.5
0
LCAT4
LCAT3
d
Relative mRNA level
2
1.5
1
0.5
0
WT
ami:lcat4
e
Relative mRNA level
1.5
1
0.5
0
WT
ami:lcat3

### Figure S6

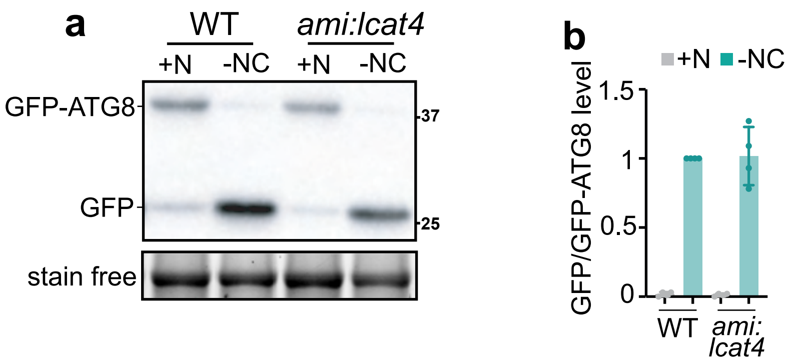
a
WT
ami:lcat4
+N
-NC
+N
-NC
GFP-ATG8
37
GFP
25
stain free
b
+N
-NC
GFP/GFP-ATG8 level
1.5
1
0.5
0
WT
ami:
lcat4

**Figure S7**

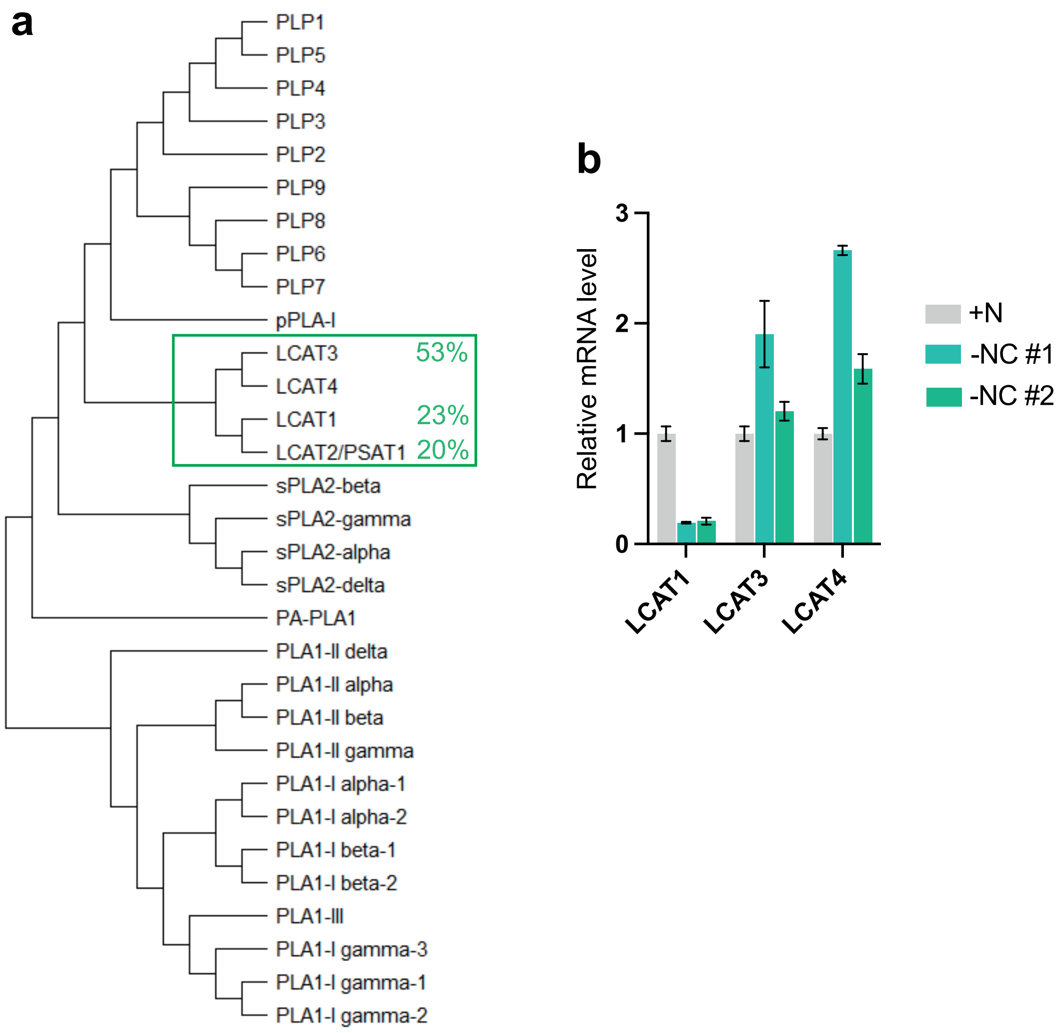
a
PLP1
PLP5
PLP4
PLP3
PLP2
PLP9
PLP8
PLP6
PLP7
pPLA-I
LCAT3
53%
LCAT4
LCAT1
23%
LCAT2/PSAT1
20%
sPLA2-beta
sPLA2-gamma
sPLA2-alpha
sPLA2-delta
PA-PLA1
PLA1-II delta
PLA1-II alpha
PLA1-II beta
PLA1-II gamma
PLA1-I alpha-1
PLA1-I alpha-2
PLA1-I beta-1
PLA1-I beta-2
PLA1-III
PLA1-I gamma-3
PLA1-I gamma-1
PLA1-I gamma-2
b
Relative mRNA level
3
2
1
0
LCAT1
LCAT3
LCAT4
+N
-NC #1
-NC #2

### Figure S8

**a**

```
LCAT4   MSLLLEEIIRSVEALLKLRNRNQEPYVDPNLNPVLLVPGIAGSILNAVDHENGNEERVWV   60
LCAT3   --------MGWIPCPCWGTNDDENAGEVADRDPVLLVSGIGGSILHSKKKNSKSEIRVWV   52

LCAT4   RIFGADHEFRTKMWSRFDPSTGKTISLDPKTSIVVPQDRAGLHAIDVLDPDMIVG---RE   117
LCAT3   RIFLANLAFKQSLWSLYNPKTGYTEPLDDNIEVLVPDDDHGLYAIDILDPSWFVKLCHLT   112

LCAT4   SVYYFHEMIVEMIGWGFEEGKTLFGFGYDFRQSNRLQETLDQFAKKLETVYKASGEKKIN   177
LCAT3   EVYHFHDMIEMLVGCGYKKGTTLFGYGYDFRQSNRIDLLILGLKKKLETAYKRSGGRKVT   172

LCAT4   VISHSMGGLLVKCFMGLHSDIFEKYVQNWIAIAAPFRGAPGYITSTLLNGMSFVNGWEQN   237
LCAT3   IISHSMGGLMVSCFMYLHPEAFSKYVNKWITIATPFQGAPGCINDSILTGVQFVEGLESF   232

LCAT4   FFVSKWSMHQLLIECPSIYELMCCPYFKWELPPVLELWREKESNDGVGTSYVVLESYCSL   297
LCAT3   FFVSRWTMHQLLVECPSIYEMMANPDFKWKKQPEIRVWRKKSENDVD--TSVELESFGLI   290

LCAT4   ESLEVFTKSLSNNTADYCGESIDLPFNWKIMEWAHKTKQVLASAKLPPKVKFYNIYGTNL   357
LCAT3   ESIDLFNDALKNNELSYGGNKIALPFNFAILDWAAKTREILNKAQLPDGVSFYNIYGVSL   350

LCAT4   ETPHSVCYGNEKMPVKDLTNLRYFQPTYICVDGDGTVPMESAMADGLEAVARVGVPGEHR   417
LCAT3   NTPFDVCYGTETSPIDDLSEICQTMPEYTYVDGDGTVPAESAAAAQFKAVASVGVSGSHR   410

LCAT4   GILNDHRVFRMLKKWLNVGEPDPFYNPVNDYVILPTTYEFEKFHENGLEVASVKESWDII   477
LCAT3   GLLRDERVFELIQQWLGVEPKKAKRKHLRTHKVVDS----------G-------------   447

LCAT4   SDDNNIGTTGSTVNSISVSQPGDDQNPQAEARATLTVQPQSDGRQHVELNAVSVSVDA   535
LCAT3   ----------------------------------------------------------   447
```

**b**

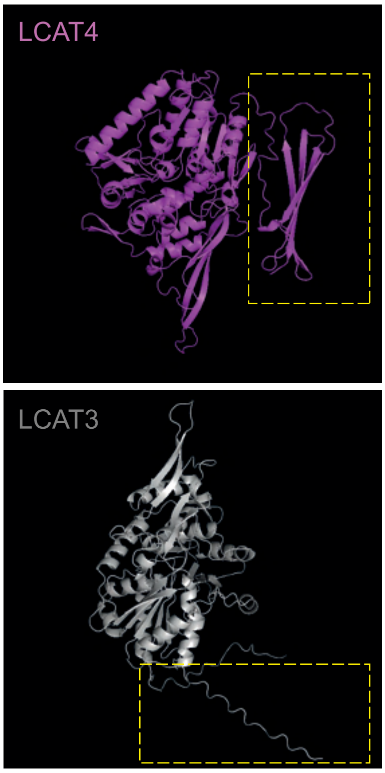


**c**

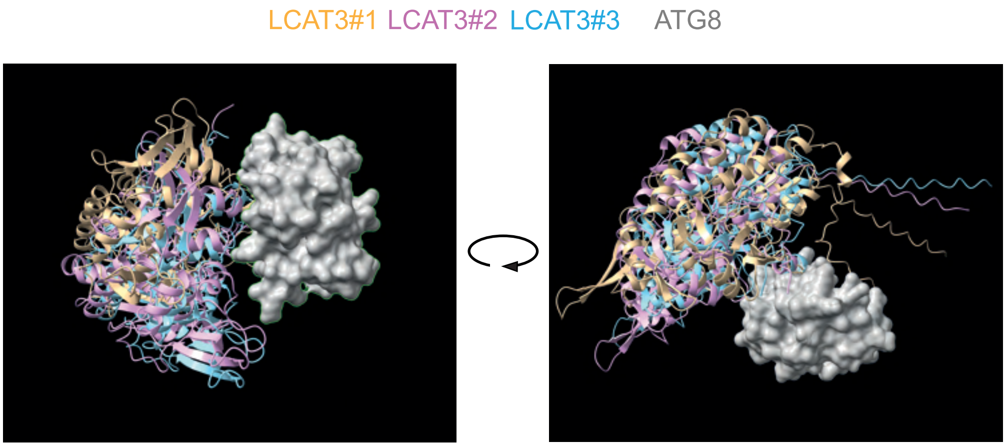


**d**

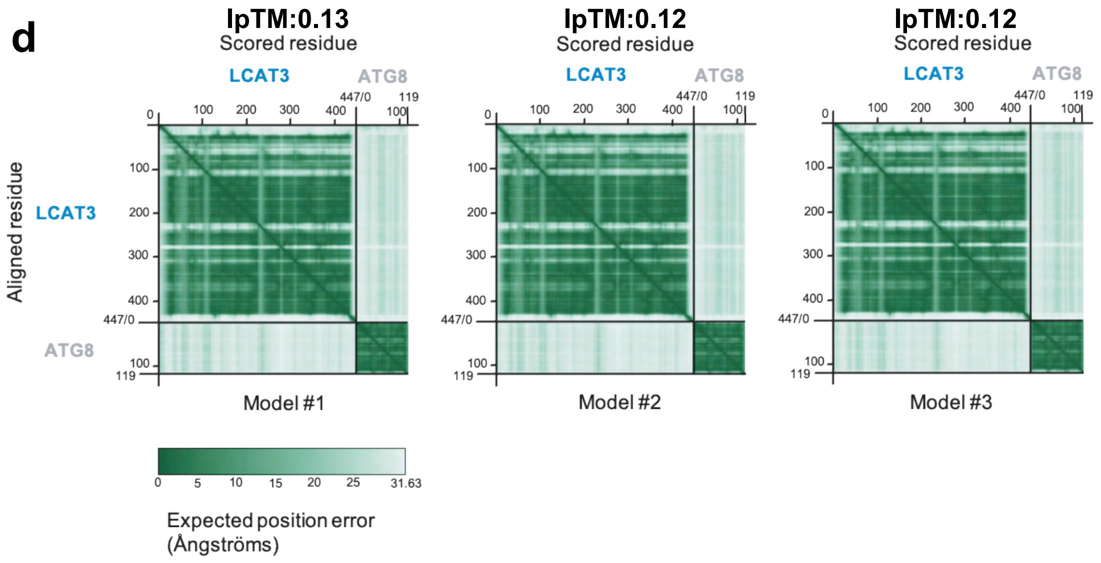

### Figure S9

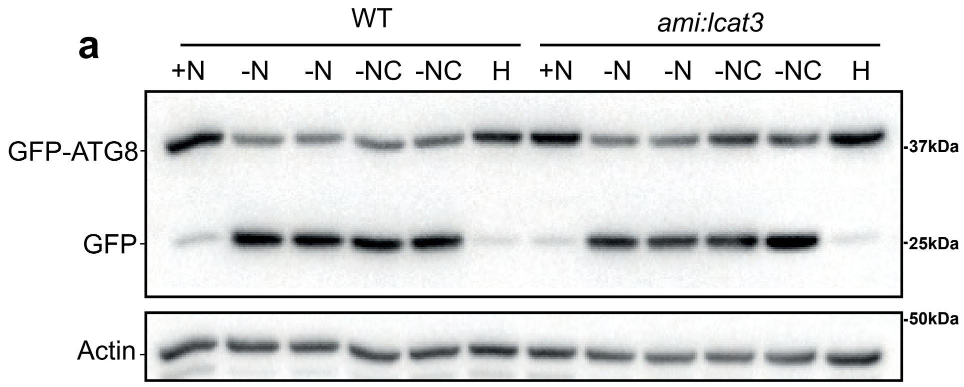

a
WT
ami:lcat3
+N -N -N -NC -NC H +N -N -N -NC -NC H
GFP-ATG8
GFP
Actin
37kDa
25kDa
50kDa


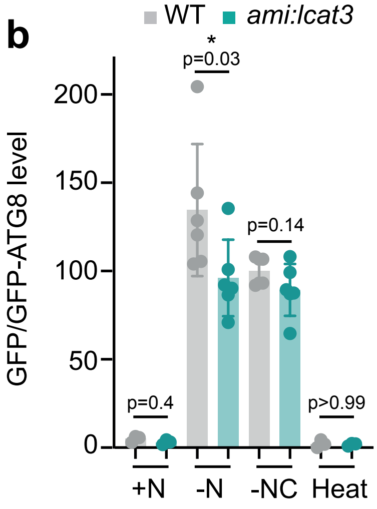

b
WT
ami:lcat3
GFP/GFP-ATG8 level
*
p=0.03
p=0.14
p=0.4
p>0.99
0
50
100
150
200
+N
-N
-NC
Heat

#### Figure S10

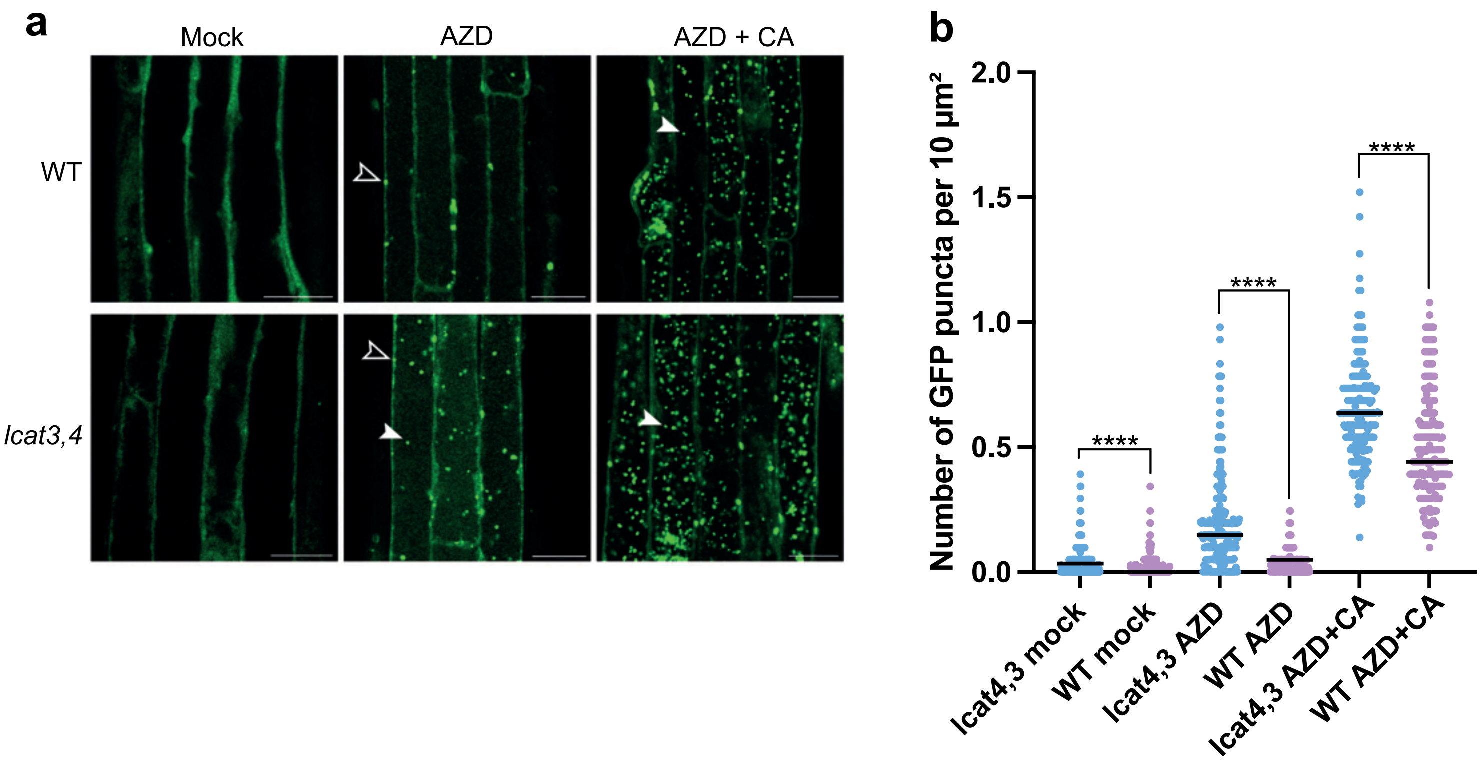
a
Mock
AZD
AZD + CA
WT
lcat3,4
b
Number of GFP puncta per 10 µm²
2.0
1.5
1.0
0.5
0.0
****
****
****
lcat4,3 mock
WT mock
lcat4,3 AZD
WT AZD
lcat4,3 AZD+CA
WT AZD+CA

#### Figure S11

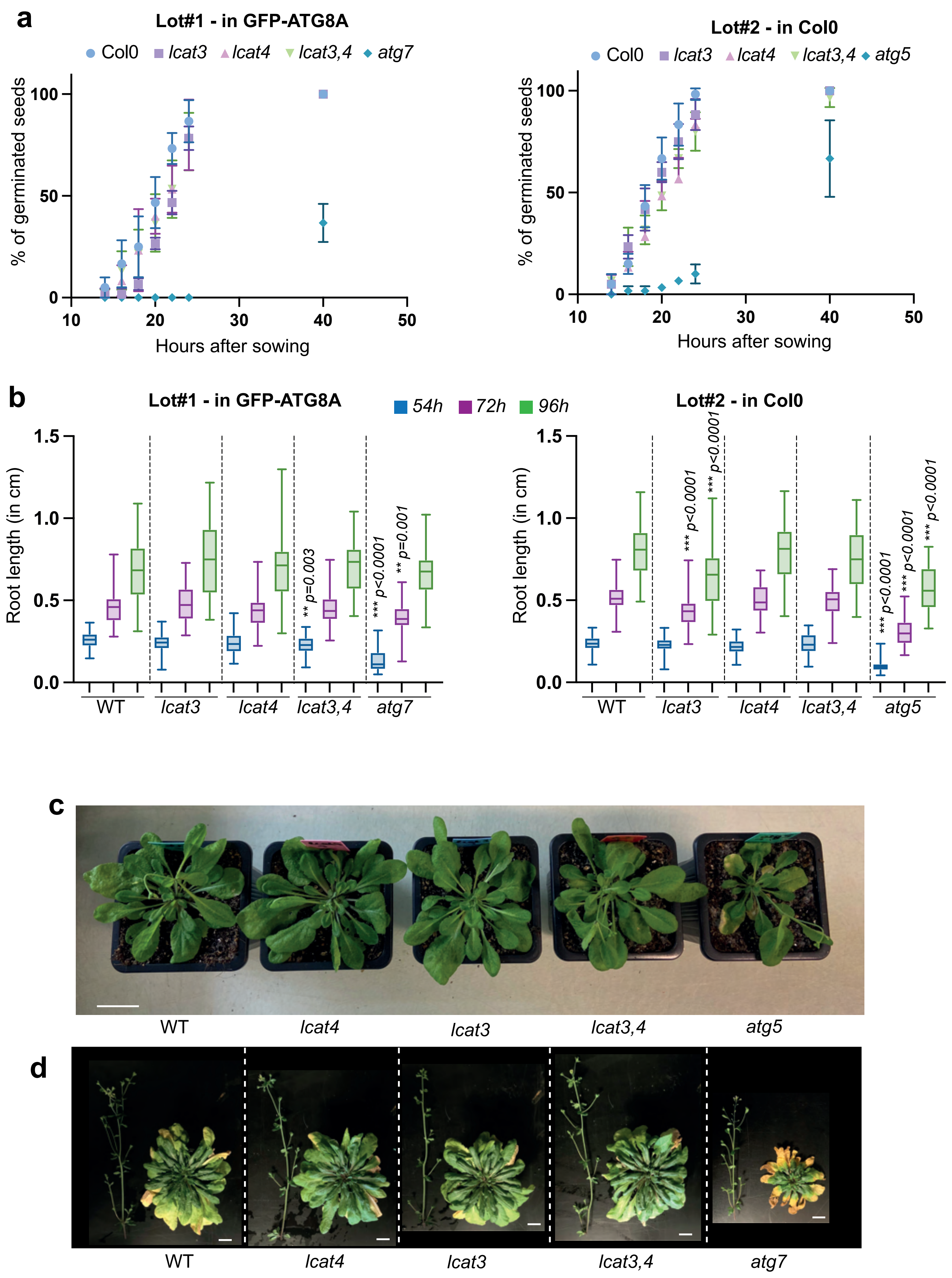

a
Lot#1 - in GFP-ATG8A
Col0
lcat3
lcat4
lcat3,4
atg7
% of germinated seeds
Hours after sowing
Lot#2 - in Col0
Col0
lcat3
lcat4
lcat3,4
atg5
% of germinated seeds
Hours after sowing
b
Lot#1 - in GFP-ATG8A
54h
72h
96h
Lot#2 - in Col0
Root length (in cm)
** p=0.003
*** p<0.0001
** p=0.001
*** p<0.0001
*** p<0.0001
*** p<0.0001
*** p<0.0001
*** p<0.0001
WT
lcat3
lcat4
lcat3,4
atg7
WT
lcat3
lcat4
lcat3,4
atg5
c
WT
lcat4
lcat3
lcat3,4
atg5
d
WT
lcat4
lcat3
lcat3,4
atg7

#### Figure S12

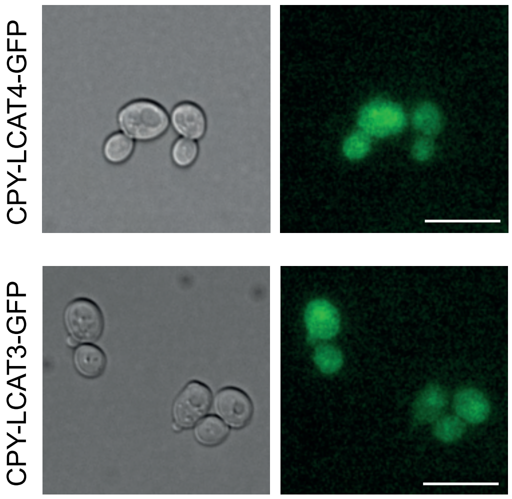
